# Evolutionary conserved *cis-trans* regulation machinery for diterpenoid phytoalexin production in Poaceae

**DOI:** 10.1101/2024.05.05.592300

**Authors:** Youming Liu, Shiho Tomiyama, Ikuya Motegi, Naoki Yamamoto, Aiping Zheng, Masaki Mori, Maki Kawahara, Yoshimasa Tsujii, Koji Miyamoto, Hiroyasu Furumi, Yutaka Sato, Hideaki Nojiri, Kazunori Okada

**Author notes:** Corresponding author:Kazunori Okada. These authors equally contributed to this work (Youming Liu, Shiho Tomiyama).

## Abstract

- Momilactones and phytocassanes are diterpenoid phytoalexins involved in plant chemical defense. These metabolites, along with biosynthetic gene clusters (BGCs), are conserved in wild rice. However, the mechanisms by which phytoalexins are regulated in wild rice are unclear. Thus, we aimed to investigate the regulatory mechanisms for biosynthetic genes within the BGCs of diterpenoid phytoalexins.
- We conducted a transcriptome analysis of five wild rice species, *Oryza rufipogon*, *Oryza punctata*, *Oryza officinalis*, *Oryza brachyantha*, and *Leersia perrieri*, after CuCl_2_ treatment.
- Among the CuCl_2_-responsive transcription factors, diterpenoid phytoalexin factor (DPF), which regulates phytoalexin production in cultivated rice (*Oryza sativa*), was broadly conserved in wild rice and showed phytoalexin-inducing activity when introduced into cultivated rice. Highly conserved genomic regions containing N-boxes (5′-CACGAG-3′), the potential binding motif of DPF, were found. CRISPR/Cas9 genome editing to remove these regions showed that biosynthetic gene expression and phytoalexin production were significantly attenuated after CuCl_2_ treatment in the leaves of the edited plants. Thus, the *cis-trans* factor combination of DPF and N-boxes is a key determinant of regulation.
- DPF has evolved as a strong *cis-trans* regulatory system for diterpenoid phytoalexin production, with N-boxes generated within the cluster region during the evolution from wild rice to cultivated rice.

## Introduction

Phytoalexins are low-molecular-weight secondary metabolites in plants that exhibit broad-spectrum antimicrobial activities (Ahuja *et al*., 2012). Biosynthetic genes of certain plant secondary metabolites, including terpenoids, gather in genomes to form biosynthetic gene clusters (BGCs). Such BGCs have been discovered in several plant species. For instance, in tobacco (*Nicotiana tabacum*), the BGCs responsible for sesquiterpenoid phytoalexin capsidiol biosynthesis have been characterised using *in silico* approaches and have been suggested to be conserved in the *Nicotiana* genus (Chen *et al*., 2019). In maize (*Zea mays*), biosynthetic genes for the sesquiterpenoid phytoalexin zealexin have been integrated as a BGC into chromosomes (Ding *et al*., 2020). Two diterpenoid phytoalexins, momilactones and phytocassanes, have been isolated from cultivated rice (*Oryza sativa*) (Cartwright *et al*., 1981; Koga *et al*., 1995, 1997; Horie *et al*., 2015). Momilactones and phytocassanes accumulate in response to different environmental stimuli, such as ultraviolet irradiation (UV), copper chloride (CuCl_2_), and pathogenic infections (Cartwright *et al*., 1981; Kodama *et al*., 1988a,b; Hasegawa *et al*., 2010). The corresponding biosynthetic genes for momilactones and phytocassanes are also involved in BGC formation on chromosomes 4 and 2 in *O. sativa* (Shimura *et al*., 2007; Wang *et al*., 2012).

BGCs associated with the production of momilactones and phytocassanes are conserved in multiple wild rice species (Miyamoto *et al*., 2016). These species encompass the AA genome species *Oryza rufipogon*, which hosts complete sets of BGCs for momilactones and phytocassanes, because of its close genetic affinity with cultivated rice. The BB genome *Oryza punctata* possesses a complete BGC for momilactones but an incomplete BGC for phytocassanes. The FF genome species *Oryza brachyantha* features only two incomplete BGCs, with some genes encoding cytochrome P450 enzymes. Finally, the outgroup species *Leersia perrieri* possesses a complete BGC of phytocassanes, which was suggested to be arranged and formed independently of the *Oryza* genus (Miyamoto *et al*., 2016). Genome information on *Oryza officinalis* has been recently released, but the capacity for phytoalexin biosynthesis and the presence of BGCs in this species remain unclear (Shenton *et al*., 2020).

The biosynthetic genes for momilactone and phytocassane production within the clusters in the cultivated rice genome have been well studied for their enzymatic functions, and the following biosynthetic route has been proposed. Within the momilactone BGC, *OsCPS4* and *OsKSL4* encode two terpene cyclases responsible for the consecutive conversion of the precursor geranylgeranyl diphosphate (GGDP), derived from the methylerythritol phosphate (MEP) pathway, into the intermediate product *syn*-pimaradiene via *syn*-copalyl diphosphate (Peters 2006; Toyomasu 2008; Okada *et al*., 2007). Subsequently, cytochrome P450 and dehydrogenase genes located inside or outside the BGC encode enzymes that catalyse the conversion of intermediate products into a final product known as momilactone A (Shimura *et al*., 2007; Wang *et al*., 2011; Toyomasu *et al*., 2014). Notably, the BGC of momilactones is incomplete and fragmented, necessitating the presence of two cytochrome *P450* genes located on chromosomes 2 (*OsCYP76M8*) and 6 (*OsCYP701A8*) (Kitaoka *et al*., 2015, 2021). Additionally, OsCYP76M14, a cytochrome P450 enzyme located on chromosome 1, is involved in the conversion of momilactone A to momilactone B (De La Peña *et al*., 2021). Within the BGC of phytocassanes, *OsCPS2* and *OsKSL7*, which encode terpene cyclases, are responsible for the two-step cyclization of GGDP to *ent*-cassa-12,15-diene. Similar to the BGC of momilactones, the BGC of phytocassanes also contains several genes encoding cytochrome P450 enzymes that finalise phytocassane biosynthesis from intermediate products (Wang *et al*., 2012; Toyomasu *et al*., 2015; Ye *et al*., 2018). Moreover, *OsCYP76M8*, which encodes an enzyme suggested to be involved in momilactone biosynthesis, was integrated into the phytocassane BGC (Kitaoka *et al*., 2021). This evidence suggests the interdependent evolution of momilactone and phytocassane BGCs.

The advantage of plant BGCs, which integrate genes responsible for specific metabolites, is that they facilitate co-evolution and co-regulation (Chu *et al*., 2011). In cucumber, two transcription factors have been proposed as direct regulators of genes within the BGC responsible for triterpenoids called cucurbitacins (Shang *et al*., 2014). In our previous study, we identified several transcription factors that serve as activators in regulating momilactone and phytocassane biosynthesis in cultivated rice. OsTGAP1 is a bZIP-type transcription factor that exhibits root-specific expression in response to stress (Okada *et al*., 2009; Miyamoto *et al*., 2014; Yoshida *et al*., 2017). In addition, diterpenoid phytoalexin factor (DPF), a basic helix-loop-helix (bHLH)-type trans-activator, plays a critical role in regulating momilactone and phytocassane biosynthesis. Co-expression analysis has shown that *DPF* expression exhibits a robust correlation with the genes responsible for momilactone and phytocassane biosynthesis. Moreover, DPF possibly plays a critical role in regulation because of its presumed capacity to interact with the N-box (5′-CACGAG-3′) *cis*-element in the promoter region of biosynthetic genes for momilactone and phytocassane production (Yamamura *et al*., 2015). In addition to OsTGAP1 and DPF, several OsWRKY transcription factors participate in this regulatory mechanism (Yokotani *et al*., 2013; Akagi *et al*., 2014; Fukushima *et al*., 2016).

Although certain wild rice species produce momilactones and phytocassans and possess BGCs that are possibly involved in their biosynthesis, the regulatory mechanisms underlying these processes remain unknown. In this study, we aimed to elucidate the mechanisms underlying the regulation of biosynthetic genes within the BGCs of diterpenoid phytoalexins. We conducted a transcriptome analysis of five wild rice species, *O. rufipogon*, *O. punctata*, *O. officinalis*, *O. brachyantha*, and *L. perrieri*, after CuCl_2_ treatment. The expression levels of several transcription factors associated with the genes within the BGCs increased after CuCl_2_ treatment, including homologues of *DPF* conserved in all wild rice species. Molecular and transgenic approaches were also employed to investigate the functions of wild rice DPF homologues. This study provides novel insights into the evolutionarily conserved *cis-trans* regulation system of biosynthetic genes within BGCs for diterpenoid phytoalexins.

## Materials and Methods

### Plant materials and growth conditions

Seeds of *O. brachyantha* (accession W1711), *O. rufipogon* (accession W1943), *O. punctata* (accession W1514), *O. officinalis* (accession W0002), and plantlet of *L. perrieri* (W1529) were obtained from the National Institute of Genetics (Shizuoka, Japan). These plants were cultivated on soil in a biotron at 28°C with a 14-hour light and 10-hour dark condition until they vigorously grew. Cultivated rice *Oryza sativa* L. ‘Nipponbare’ was used as the protoplast material for transient assays. The seeds were sterilised with 75% ethanol for 1 min and then rinsed with tap water. Subsequently, the sterilised seeds were incubated in sterile soil with an 8 h light and 16 h dark photoperiod at 28°C for 2 weeks.

For transgenic rice plants *OsKSL4pΔ* and *OsCPS2pΔ*, sterilised seeds prepared by the above-mentioned procedure were germinated in sterilised soil with a 16 h light and 8 h dark photoperiod at 28°C.

### Chemical treatment

Approximately 3–5 expanded leaves were collected from each wild rice plant and mixed to reduce bias due to differences in the responses of independent leaves. For stress treatment, 1 cm^2^ detached leaf blades from the mature plants were soaked in 0.5 mM CuCl_2_ solution at 28°C under light for 6– 72 h. Mock (untreated) leaf blades were used as controls. These tissues were immediately frozen in liquid nitrogen and stored at -80°C until use for RNA extraction and metabolite analysis.

For *OsKSL4pΔ* and *OsCPS2pΔ*, 1-month leaves were subjected to the same treatment procedure for RNA extraction and metabolite analysis.

### mRNA Sequencing

Leaf blades were collected from the wild rice plants before (0 h control) or after CuCl_2_ treatment for 24 h, and total RNAs were extracted from powdered leaf samples using an RNeasy Plant Mini Kit (QIAGEN, Germany) in accordance with the manufacturer’s instructions. Then, the RNA fractions were incubated with RNase-free DNase I at 37°C for 30 min to remove genomic DNA contaminants. RNA integrity was verified using an Agilent Bioanalyzer 2000 (Agilent Technologies, Inc., Santa Clara, CA, USA) to confirm that the RIN values were no less than 8.0. The purity and quantity of the total RNAs were determined using a NanoDrop light spectrophotometer (Thermo Scientific, Inc., MA, USA).

For each sample, 500 ng of the extracted total RNA was used for library construction using the TruSeq RNA Sample Preparation Kit V2 (Illumina, San Diego, CA, USA) in accordance with the manufacturer’s instructions. Briefly, poly (A)^+^ RNAs were purified and fragmented for use in double-stranded cDNA synthesis. The cDNAs were ligated with an indexed adaptor and amplified by polymerase chain reaction (PCR). The resulting PCR products were purified using AMPureXP (Beckman Coulter, Brea, CA, USA). The average length of the cDNA inserts ranged from 314 bp to 329 bp. The PCR products were sequenced using HiSeq2500 (Illumina) to generate 100 bp single-end raw reads. The raw sequence data were deposited in the NCBI Sequencing Read Archive (PRJNA1096438).

Low-quality sequences and adaptor sequences on the raw Illumina reads were trimmed by Trimmomatic version 0.33 (Bolger *et al*., 2014) with the options of ‘LEADING:15, TRAINLING:15, SLIDINGWINDOW:4:15, MINLEN:75’ to prepare high-quality reads. The high-quality reads were mapped to the corresponding reference genome sequences: *O. rufipogon*, *O. punctata*, *O. brachyantha*, and *L. perrieri* of EnsemblPlants release 44 (Yates *et al*., 2022) and *O. officinalis* (provided by Shenton *et al*. 2020 before publication) using STAR version 2.7.2b (Dobin *et al*., 2013). The number of reads per gene was counted. Read count data were normalised using TCC (Sun *et al*., 2013) and converted to transcripts per million (TPM) (Wagner *et al*., 2012). Differentially expressed genes (DEGs) were identified using edgeR with a generalised linear model at the 5% level of statistical significance (Robinson *et al*., 2010). Probable orthologous genes between *O. sativa* and the wild rice species were predicted based on BLASTP searches, with an E-value of 1e^-10^. BGCs’ gene annotations for *O. officinalis* were based on the corresponding *O. sativa* genes. Gene annotations for rice transcription factors in *O. sativa* were retrieved from PlantTFDB ver. 5.0 (Guo et al., 2008). Gene set enrichment analyses of DEGs were conducted using the web-based rice omics tool CARMO (Wang et al., 2015). Under the assurance that the gene populations in *O. sativa* and the wild rice species analysed were not different, GO and gene module enrichment analyses were performed. Identifiers of the predicted orthologous *O. sativa* genes to DEGs in wild rice were queried using this web tool to obtain enrichment test results. We applied a statistically significant false discovery rate (FDR) threshold of less than 0.05.

### Expression analysis

Leaf blades from the wild rice plants were collected after 6, 12, 24, 48, and 72 h of CuCl_2_ treatment. Leaf blades from the wild-type (WT) and transgenic rice plants *OsKSL4pΔ* and *OsCPS2pΔ* were collected after 6 h of CuCl_2_ treatment to investigate the expression levels of *OsCPS2* and *OsKSL4*. Total RNA was extracted by Sepasol^®^-RNA I Super G (Nacalai tesque, Kyoto, Japan). cDNA was reverse transcribed using the PrimeScript^TM^ RT reagent kit (Perfect Real Time) (TaKaRa, Shiga, Japan). The cDNA mixture was used for qRT-PCR using GoTaq^®^ qPCR and RT-qPCR Systems (Promega, WI, USA) on a 7300 Real-Time PCR System (Applied Biosystems). The 2^–△△CT^ method was used to calculate relative expression levels (Livak *et al*., 2001).

### Plasmid construction and plant transformation

The *DPF* open-reading frame fragments from the wild rice species were PCR amplified and subsequently cloned into the pENTR/D-TOPO vector (Invitrogen, MA, USA). The fragments were then inserted into the transient expression vector pUbi-Rfa-Tnos using the LR reaction (Gateway^®^). pUbi-*OsDPF*-Tnos, pUbi-*OrDPF*-Tnos, pUbi-*OpDPF*-Tnos, pUbi-*OoDPF*-Tnos, pUbi-*ObDPF*-Tnos, and pUbi-*LpDPF*-Tnos were constructed, and their expression was controlled by the maize polyubiquitin promoter. pUbi-*GUS*-Tnos (Chujo *et al*., 2014), in which the expression of *β-*glucuronidase (GUS) was controlled by the polyubiquitin promoter, served as a negative control.

For the reporter gene assay, pUbi-*RLUC*-Tnos (Ogawa *et al*., 2017) expressing *renilla luciferase* was used as the internal control. pGL3-*CYP99A2p*-2.4kb, pGL4.1-*CPS2p*-0.4kb, and pGL3-*KSL4p*-2.0kb, in which the promoter regions of *OsCYP99A2*, *OsCPS2*, and *OsKSL4* in *O. sativa* were fused to the *firefly luciferase* (*FLUC*) gene individually, were used as reporter vectors.

For CRISPR/Cas9 construction, the guide RNA cloning vectors pU6gRNA and pZH_gYSA_MMCas9 were used (Mikami *et al.,* 2015). Then, 20 bp sequences from -332 to -352 and -463 to -483 of the *OsCPS2* promoter region and from -1003 to -1023 and -1962 to -1982 of the *OsKSL4* promoter region (including PAM) were selected as the target sites of Cas9 by using the guide RNA design tool CRISPR-P 2.0 (Liu et al., 2017). The DNA fragments were cloned into pU6gRNA using the restriction enzyme *Bbs*I. These DNA fragments were subcloned into pZH_gYSA_MMCas9. The CRISPR/Cas9 plasmids were transformed into *Agrobacterium tumefaciens* strain EHA105 to transform calli generated from rice embryos. The genomic regions containing the Cas9 target sites were PCR amplified to confirm the mutations.

### Protoplast isolation and transient assay

The method of protoplast isolation was primarily based on the protocol described by Zhang *et al*. (2011). In brief, the sheath portion of 2-week rice seedlings was dissected into strips, submerged in an enzymatic digestion buffer [1% Cellulase Onozuka^TM^ R-10 (Yakult Pharmaceutical, Tokyo, Japan), 0.5% Macerozyme^TM^ R-10 (Yakult Pharmaceutical, Tokyo, Japan), 0.1% bovine serum albumin (Sigma-Aldrich, MO, USA), 0.6 M mannitol, 10 mM 2-(N-morpholino)ethanesulphonic acid (MES) at pH 5.7, and 10 mM CaCl_2_], and then incubated for 4 h in the dark with gentle shaking. After enzymatic digestion, the residue was filtered, and protoplasts were isolated using W5 buffer (154 mM NaCl, 125 mM CaCl_2_, 5 mM KCl, and 2 mM MES at pH 5.7).

In the transient phytoalexin measurement, polyethylene glycol (PEG)-mediated transformation was performed for plasmid introduction. For each sample, 10 μg of plasmid, 100 μl of protoplast solution (2 × 10^5^ cells), 110 μl of 40% PEG 4000 solution (Sigma-Aldrich), 0.2 M mannitol, and 0.1 M CaCl_2_ were mixed and incubated in the dark for 15 min. Following transformation, the samples were incubated in W5 buffer and stored in the dark at 25°C for 2 days.

For the reporter gene assay, pUbi plasmids containing *DPF* homologues or *GUS* were used as effectors. pGL3-*CYP99A2p*-2.4kb, pGL4.1-*CPS2p*-0.4kb, and pGL3-*KSL4p*-2.0kb were used as reporters. pUbi-*RLUC*-Tnos served as the internal control. In brief, 5 μg of effector, 5 μg of reporter, and 1 μg of the internal control were transformed into 2.0 × 10^5^ cells using the PEG-mediated method described above. Following transformation, the samples were incubated in W5 buffer and stored overnight in the dark at 25°C. Subsequently, the samples were collected, and the signal intensities of LUC and RLUC were quantified using the Dual-Luciferase^®^ Reporter Assay System (Promega) on GloMax^®^ 20/20 Luminometer (Promega).

### Diterpenoid phytoalexin measurement

The CuCl_2_-treated leaf blades from the wild rice species were submerged in 2 ml of 80% methanol at 4°C for 24 h, and 5 μl of the extract was subjected to LC-MS/MS with the reaction monitoring transitions used as previously described (Miyamoto *et al*., 2016). For transient phytoalexin measurements, phytoalexin extraction was carried out using 100% methanol at 4°C for 6 h. The supernatant (5 µl) from the extracted samples were used for phytoalexin measurement.

### Statistical analysis

Statistical analyses were performed using R ver. 4.1.2. The significance of the differences between the control and experimental groups was determined using one-tailed Student’s *t*-test.

## Results

### Inductive accumulation of diterpenoid phytoalexin in wild rice species

In our previous study, we assessed the abilities of *O. rufipogon*, *O. punctata*, *O. brachyantha* and *L. perrieri* to produce momilactones and phytocassanes (Miyamoto *et al*., 2016). In the present study, we characterised the BGCs responsible for phytoalexin production in the recently sequenced species *O. officinalis* (Shenton *et al*., 2021). When searching for homologues of *O. sativa* genes, including cyclase genes *OsCPS4* and *OsKSL4*, cytochrome *P450* genes *OsCYP99A2* and *OsCYP99A3*, and dehydrogenase momilactone A synthase (*OsMAS*) within the genome assembly of *O. officinalis*, we identified a potential momilactone BGC that comprises five cyclases (three *CPS4s* and two *KSL4*s, including pseudogenes), two cytochrome *P450* genes, and an *MAS* gene located within a 400 kb region on chromosome 4 (Fig. **1a**). This discovery represents the longest momilactone BGC found to date and marks the first instance of *CPS4* and *KSL4* duplications within experimentally functional momilactone BGCs (Priego Cubero *et al*., 2024). Regarding potential phytocassane biosynthetic genes, we identified homologues of *O. sativa* genes, including *OsCPS2*, *OsKSL5*, *OsKSL6*, *OsKSL7*, *OsKSL12*, *OsKSL13*, and cytochrome *P450* genes *OsCYP76M5-8* and *OsCYP71Z6-7,* within a potential BGC located in a 500 kb region on chromosome 2 (Fig. **1b**). Duplication of *KSL7* was also observed in chromosome 2 BGCs. Considering that the genome of *O. officinalis* is 1.6 times larger than that of *O. sativa*, we can reasonably assume that this species possesses longer BGCs.

**Figure 1:**
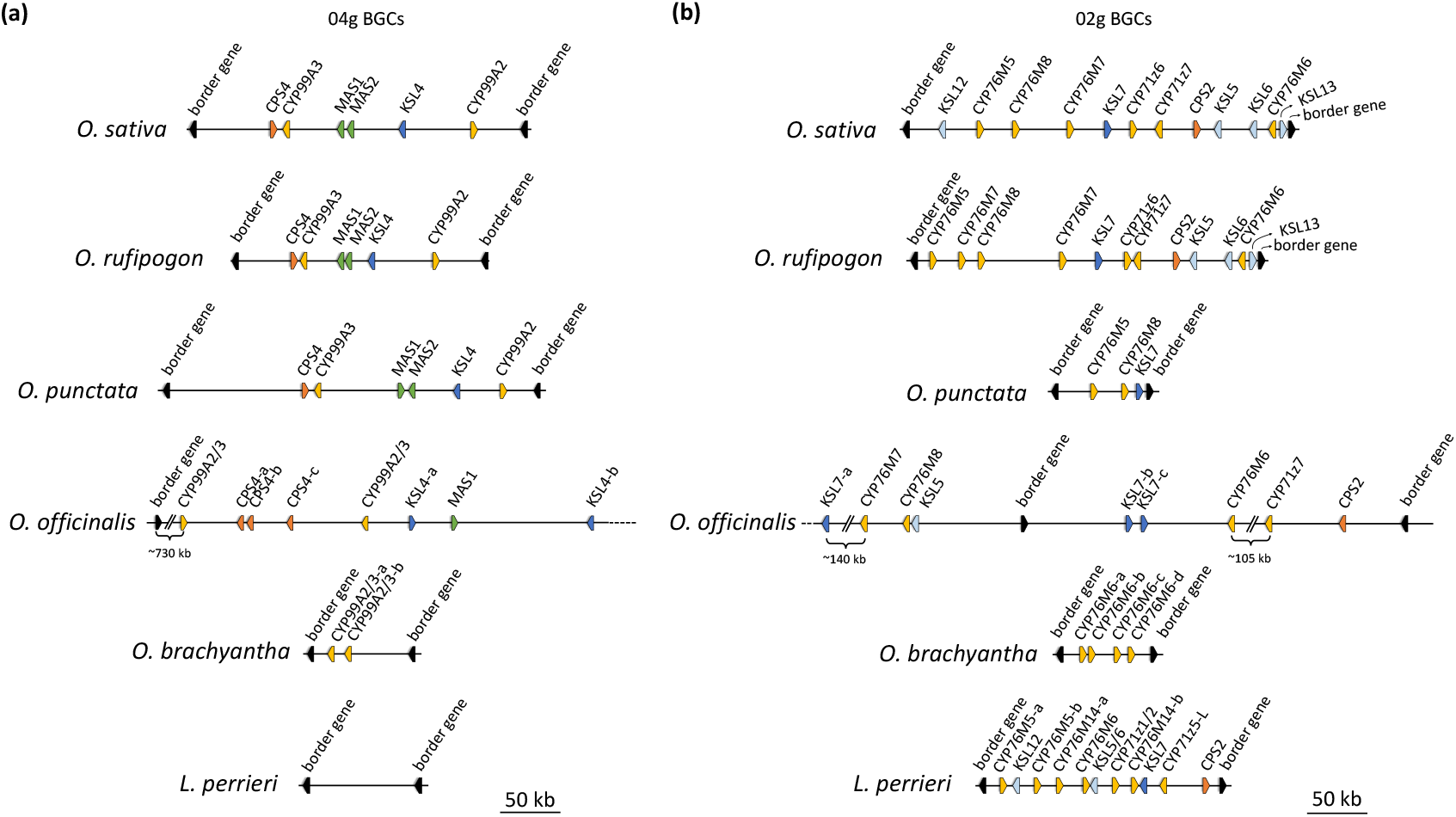
Schematic of gene loci in cultivated rice *Oryza sativa* and wild rice biosynthetic gene clusters (BGCs) on correspondent chromosomes 2 and 4. **(a)** Biosynthetic gene cluster of momilactone on chromosome 4. **(b)** BGCs of phytocassanes on chromosome 2. Details of biosynthetic gene symbol and corresponding locus IDs in *O. sativa*, *O. punctata*, *O. brachyantha*, and *L. perrieri* can be found in Miyamoto *et al*. (2016), and details of *O. officinalis* genes can be found in Table S1.

To further investigate the inducible profiles of these wild rice species, we treated the leaves of five wild rice plants with CuCl_2_. Initially, we verified the inducible profiles of cyclase and cytochrome *P450* genes in BGCs. The expression of key biosynthetic genes in the BGCs (*OrKSL7*, *OrKSL4*, *OpKSL4*, *OpCYP76M8-a*, *OoCPS2*, *OoCPS4-c*, *ObCYP76M6*, *ObCYP99A2/3*, and *LpKSL7*) was induced after 12 h of CuCl_2_ treatment, with a transcriptional peak after 48 h of treatment (Fig. **2a**).

**Figure 2:**
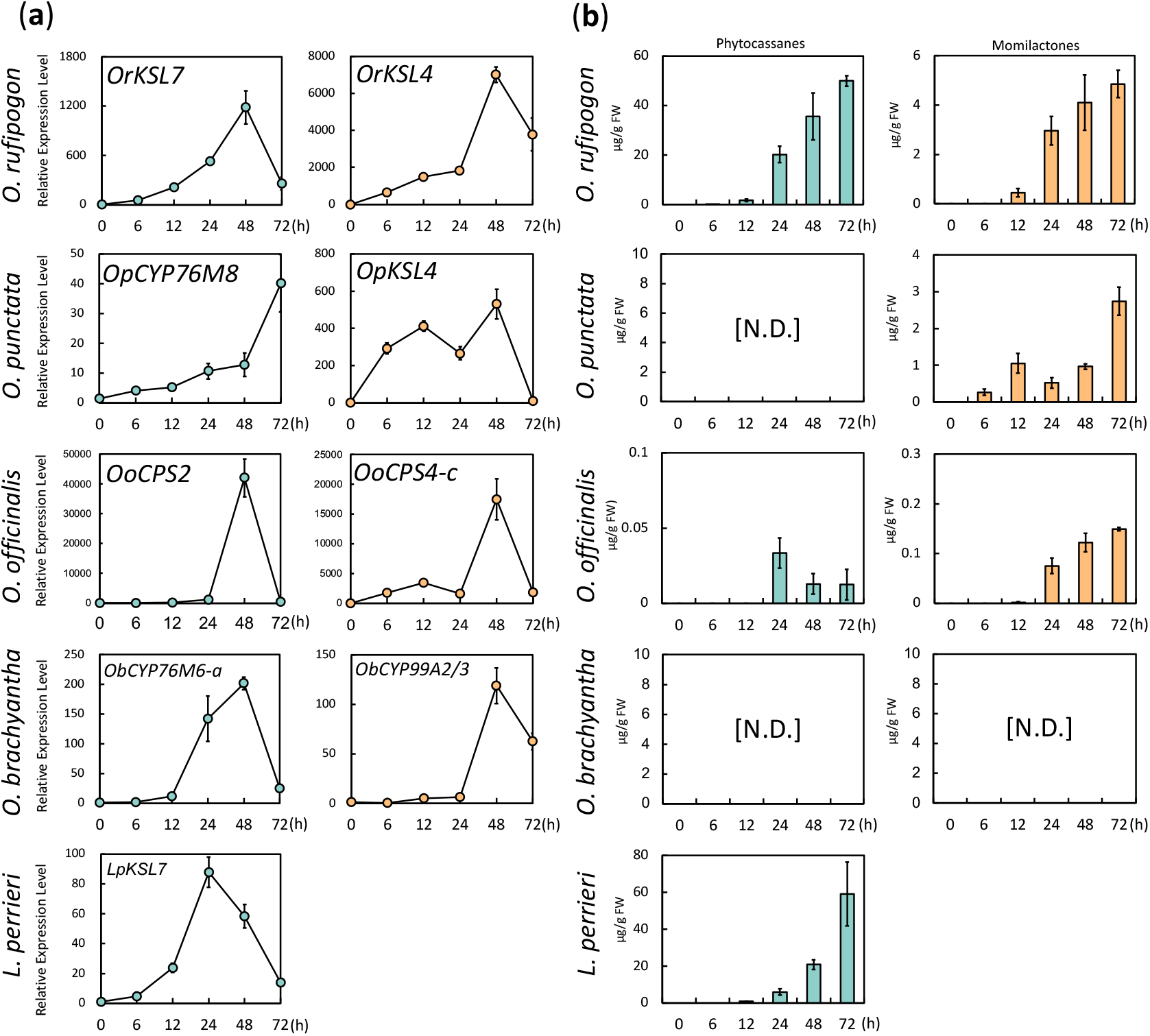
Inductive profiles of diterpenoid phytoalexins at the transcription and metabolite levels. (**a**) Expression profiles of biosynthetic genes within biosynthetic gene clusters (BGCs) after a time-course CuCl_2_ treatment (means ± SE, n = 3). *OrKSL7* and *OrKSL4* in *Oryza rufipogon*, *OpCYP76M8* and *OpKSL4* in *Oryza punctata*, *OoCPS2* and *OoCPS4-c* in *Oryza officinalis*, *ObCYP76M6-a* and *ObCYP99A2/3-a* in *Oryza brachyantha*, and *LpKSL7* in *Leersia perrieri* were selected as representative biosynthetic genes. Cyan and orange dots represent biosynthetic genes on chromosomes 2 and 4 BGCs, respectively. (**b**) Accumulation profiles of momilactones and phytocassanes after the time-course treatment with CuCl_2_. The cyan- and orange-filled bars represent the accumulation levels of phytocassanes and momilactones, respectively. N.D. indicates that a particular compound was not detected (means ± SE, n = 3).

Subsequently, we quantified the accumulation of momilactones and phytocassanes over the same time course after CuCl_2_ treatment (Fig. **2b**). These diterpenoid phytoalexins gradually accumulated after CuCl_2_ treatment in the species possessing a complete set of biosynthetic genes in their BGCs. In *O. officinalis*, we detected low but inductive momilactone and phytocassane production. Additionally, the presence of transcriptionally inducible cytochrome *P450* genes in the incomplete BGCs of *O. brachyantha* implies that this species can biosynthesise novel metabolites other than momilactones and phytocassanes. Taken together, these results suggest that the biosynthetic genes of the five wild rice species exhibit transcriptional responsiveness to stress treatments and emphasise that the capacity for diterpenoid phytoalexin production is closely linked to the presence of corresponding intact BGCs.

### Transcriptome analysis of five wild rice species under CuCl_2_ treatment

The discovery of BGCs in the CC genome species *O. officinalis* enhanced our understanding of the evolutionary trajectory of diterpenoid phytoalexin BGCs within the *Oryza* genus. Treatment with CuCl_2_ induced the expression of biosynthetic genes, accompanied by the accumulation of diterpenoid phytoalexins. This result suggests that the five wild rice species employ similar machinery to regulate their biosynthetic genes. To explore the regulatory machinery governing diterpenoid phytoalexin biosynthesis in the five wild rice species, we conducted a transcriptome analysis following CuCl_2_ treatment. In the RNA-Seq analysis, leaf tissue samples from wild rice plants treated with CuCl_2_ for 24 h served as the treatment group, whereas those just before the treatment served as the control group. Initially, we focused on the expression patterns of all biosynthetic genes located within the BGCs. As demonstrated by the heatmap in Fig. **3a**, the expression levels of most biosynthetic genes obviously increased after CuCl_2_ treatment, which corresponded with the results of our time-course experiment (Fig. **2a**), suggesting that they were regulated by a common regulatory mechanism.

**Figure 3:**
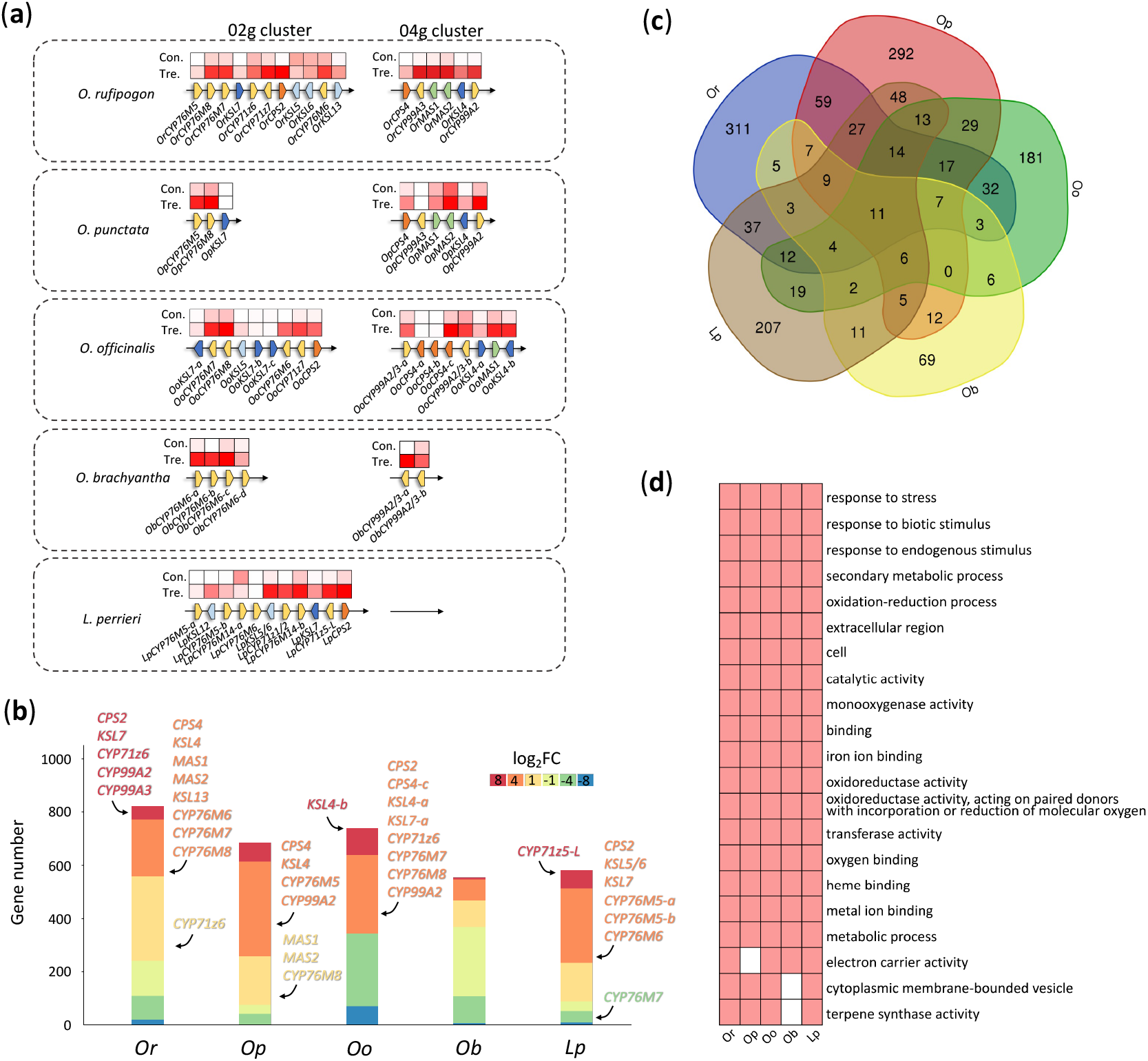
Overview of RNA-Seq analysis on deferentially expressed genes (DEGs). (**a**) Expression of biosynthetic genes within biosynthetic gene clusters. Heatmaps represent expression profiles in the control and treatment groups. Transcripts per million (TPM) (log_2_ scale) is shown in heatmaps. Red boxes represent a high expression level, whereas white boxes represent a low expression level. Con.: control group, Tre.: treatment group. (**b**) DEGs among five wild rice species. The x-axis represents the species, and the y-axis represents the number of DEGs in each species. The red and blue regions represent up-regulated and down-regulated DEGs, respectively. Deeper color indicates higher or lower fold change in up- or down-regulated DEGs, respectively. *Or*: *Oryza rufipogon*, *Op*: *Oryza punctata*, *Oo*: *Oryza officinalis*, *Ob*: *Oryza branchyantha*, *Lp*: *Leersia perrieri*. (**c**) Common DEGs across five wild rice species. (**d**) Gene Ontology enrichment analysis across five wild rice species.

We investigated the expression levels (in TPM) of all genes in the individual wild rice species. DEGs were defined based on the following criteria: log_2_FC > 1 or < -1 and FDR < 0.05. Fig. **3b** displays the number of DEGs in each wild rice species, with the colour scale indicating the fold change of these DEGs. The numbers of up-regulated and down-regulated genes varied among the different wild rice species (Table 1). *O. bracyantha* exhibited the least number of up-regulated genes (185), whereas *O. punctata* exhibited the greatest number of up-regulated genes (611). *O. rufipogon*, *O. bracyantha,* and *O. officinalis* had relatively greater numbers of down-regulated genes (240, 367, and 342, respectively) than *O. punctata* and *L. pertieri*, which showed less numbers of down-regulated genes (73 and 86, respectively). Diterpenoid phytoalexin-related biosynthetic genes were included within the gene group, most of which were placed at high fold-change values, ranging from a log_2_FC of 2–10 (Fig. **3c**).

**Table 1.** Number of differentially expressed genes (DEGs) among five wild rice species treated with 500 μM CuCl_2_ compared to mock control before the treatment.

|  | Up-regulated DEGs | Down-regulated DEGs |
| --- | --- | --- |
| *Oryza rufipogon* | 582 | 240 |
| *Oryza punctata* | 611 | 73 |
| *Oryza officinalis* | 396 | 342 |
| *Oryza bracyantha* | 185 | 367 |
| *Leersia perrieri* | 494 | 86 |

We constructed a Venn graph of all DEGs from the five wild rice varieties to identify common genes responsive to CuCl_2_ treatment. Fig. **3c** shows that 11 up-regulated genes were common to all five species, and *DXS*, which encodes deoxyxylulose phosphate synthase in the MEP pathway, was the only gene related to the production of diterpenoid phytoalexins (Dataset S1). No common down-regulated genes were found among the five species. Among the momilactone producers (*O. rufipogon*, *O. punctata*, and *O. officinalis*) or phytocassane producers (*O. rufipogon*, *O. officinalis*, and *L. perrieri*), 41 and 49 common up-regulated genes were found, respectively. They included several momilactone and phytocassane biosynthetic gene homologues *CPS2*, *CPS4*, *KSL4*, and *KSL7* in those of wild rice, together with the bHLH transcription factor DPF, a diterpenoid-regulating factor, and the NAC factor, which are thought to function in the scavenging of reactive oxygen species as commonly existing DEGs (Wu *et al*., 2022).

Gene Ontology (GO) enrichment analysis of the DEGs revealed 22 overrepresented GO terms that were shared across at least four species (Fig. **3d**). The GO terms from the up-regulated DEGs included ‘response to stress’ [GO:0006950], ‘oxidation-reduction process’ [GO:0055114], and ‘terpene synthase activity’ [GO:0010333], which are associated with stress response, oxidation/reduction, and terpenoid metabolism (Dataset S2). Gene module enrichment analysis using CARMO (Wang *et al*., 2015) clarified the over-representation of up-regulated genes in module_214, module_688, and module_748, which are involved in diterpenoid phytoalexin biosynthesis (Dataset S3). Except for the metabolic process [GO:0008152], no common over-represented GO terms from the down-regulated DEGs were found. These observations indicate that a specific gene regulation associated with CuCl_2_ response and diterpenoid phytoalexin production is maintained among the five species. We observed 59 GO terms that were overrepresented in only one species. These GO terms include ‘electron transport chain’ [GO:0022900], ‘response to heat’ [GO:0009408], and ‘response to extracellular stimulus’ [GO:0009991]. Because most of these GO terms were not related to diterpenoid phytoalexin biosynthesis, the genes assigned to these GO terms may be related to different CuCl_2_ responses among the species.

### Search for potential transcription factors involved in the regulation of phytoalexin biosynthesis

To identify the common regulatory machinery for diterpenoid phytoalexin production in the *Oryza* genus, we mined transcription factor genes that commonly respond to CuCl_2_ stress in our RNA-Seq profiles. Considering that the abovementioned common DEGs among the five species contained no orthologous transcription factor genes, we checked the transcription factor genes that were up-regulated or down-regulated in at least three species. Twenty putative orthologous gene sets were retained for further screening (Dataset S4). After manual evaluation of the fold-change status of each gene in the five species, we selected 10 potential candidate transcription factor genes: two NAC, one B3, two bHLH, one C2H2, and four WRKY. All of these genes were up-regulated after CuCl_2_ treatment, with no inconsistency (Fig. **4a**). Of these transcription factor genes, *DPF*, *WRKY27*, *WRKY32*, *AUX3*, *ONAC096*, and *RAV3* were highlighted based on their molecular conservation across the five genomes. Finally, considering the existing knowledge on diterpenoid phytoalexin production and immune response regulators (Yamamura *et al*., 2015; Abdullah-Zawawi *et al*., 2021; Wang *et al*., 2021), we selected DPF, the two WRKYs, and ONAC096 as the final candidates. No report was available on the involvement of these candidate transcription factors, except for DPF, in diterpenoid phytoalexin production. Thus, we focused on DPF homologues. Time course analysis revealed that the transcriptional profiles of each DPF homologue in wild rice inductively increased up to 48 h after CuCl_2_ treatment. In addition, *OpDPF* and *ObDPF* responded earlier to the treatment within 12 h, with biphasic expression at 48 h (Fig. **4b**).

**Figure 4:**
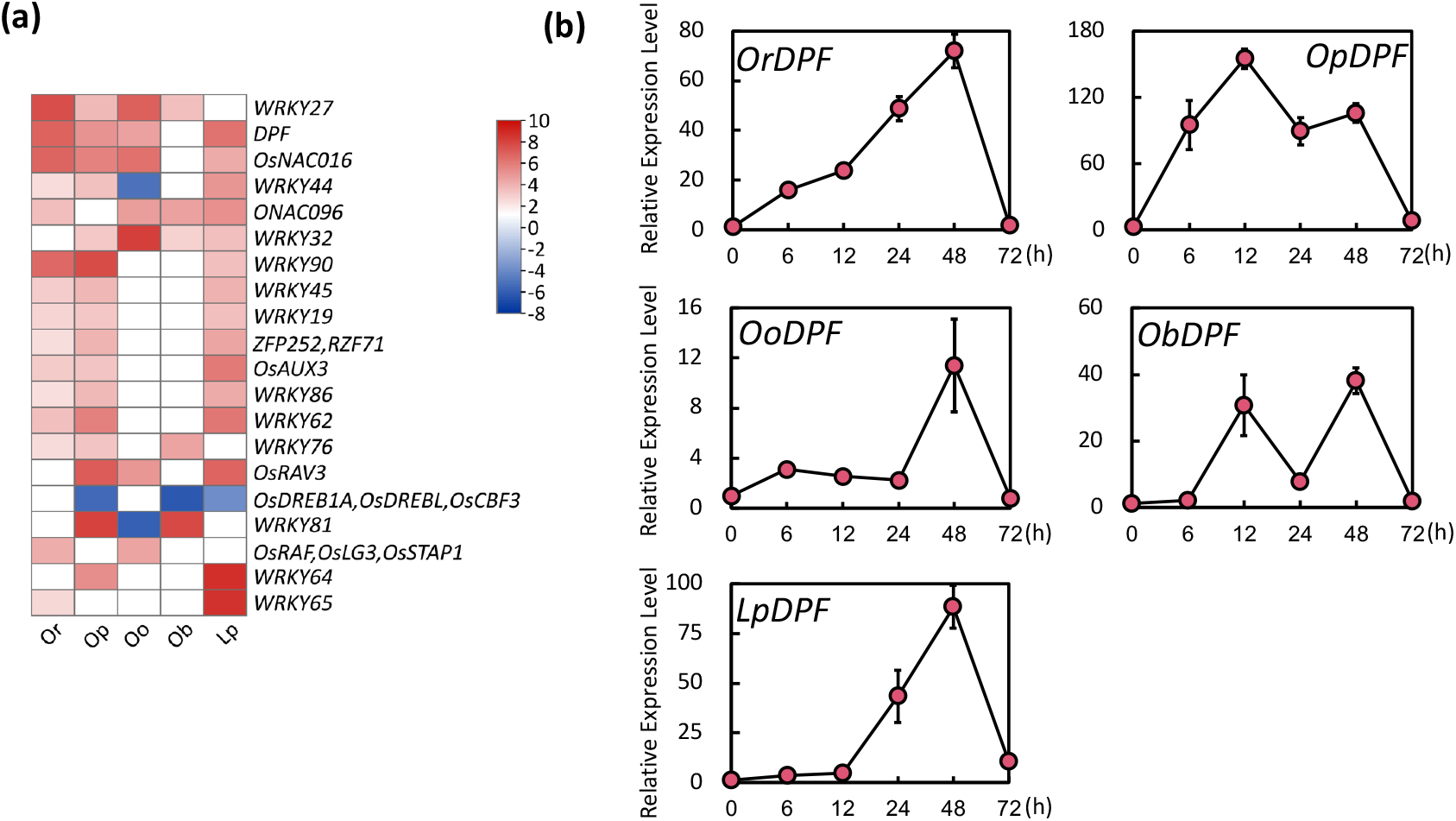
Transcription factors inductively expressed by CuCl2 treatment in five wild rice species. **(a)** List of transcription factors that were transcriptionally enhanced after CuCl_2_ treatment. Homologues of *Oryza sativa* transcription factor in wild rice species were searched using PlantTFDB V5 (http://planttfdb.gao-lab.org/). Each box represents the homolog of *O. sativa* transcription factor in a specific wild rice species, and boxes were filled with red if the fold change in RNA-Seq analysis was larger than 2. **(b)** Expression profiles of *DPF* homologues in wild rice species at different time points after CuCl_2_ treatment (means ± SE, n = 3). *OrDPF: DPF* homolog in *Oryza rufipogon. OpDPF: DPF* homolog in *Oryza punctata. OoDPF: DPF* homolog in *Oryza officinalis. ObDPF: DPF* homolog in *Oryza brachyantha. LpDPF: DPF* homolog in *Leersia perrieri*.

### Functional analysis of *DPF* homologues in wild rice species

All *DPF* homologues in the wild rice species exhibited transcriptional induction, suggesting their pivotal roles in the biosynthesis of diterpenoid phytoalexins. The cDNAs of the *DPF* homologues were successfully amplified through RT-PCR using total RNA from CuCl_2_-treated leaf tissues and cloned into *pUbi-Rfa-Tnos*, a transient expression vector driven by the maize ubiquitin promoter (Fig. 5a). The sequences of the *DPF* homologues in *O. punctata* and *O. brachyantha* were not correctly predicted, and the correct mRNA sequences were deduced based on the RNA-Seq mapping results (Dataset S5). Transient expression vectors containing the *DPF* homologues were individually introduced into *O. sativa* sheath protoplasts to confirm their ability to initiate diterpenoid phytoalexin production. Following incubation, diterpenoid phytoalexins were extracted from the protoplasts harbouring the vectors using methanol, and the resulting extract was subjected to LC-MS/MS analysis. Similar to *O. sativa DPF*, all DPF homologues from wild rice induced the biosynthesis of momilactones and phytocassanes (Fig. 5b). This result indicates that DPF homologues function as crucial nodes in the biosynthesis of diterpenoid phytoalexins in *Oryza* and other grass species.

**Fig. 5.**
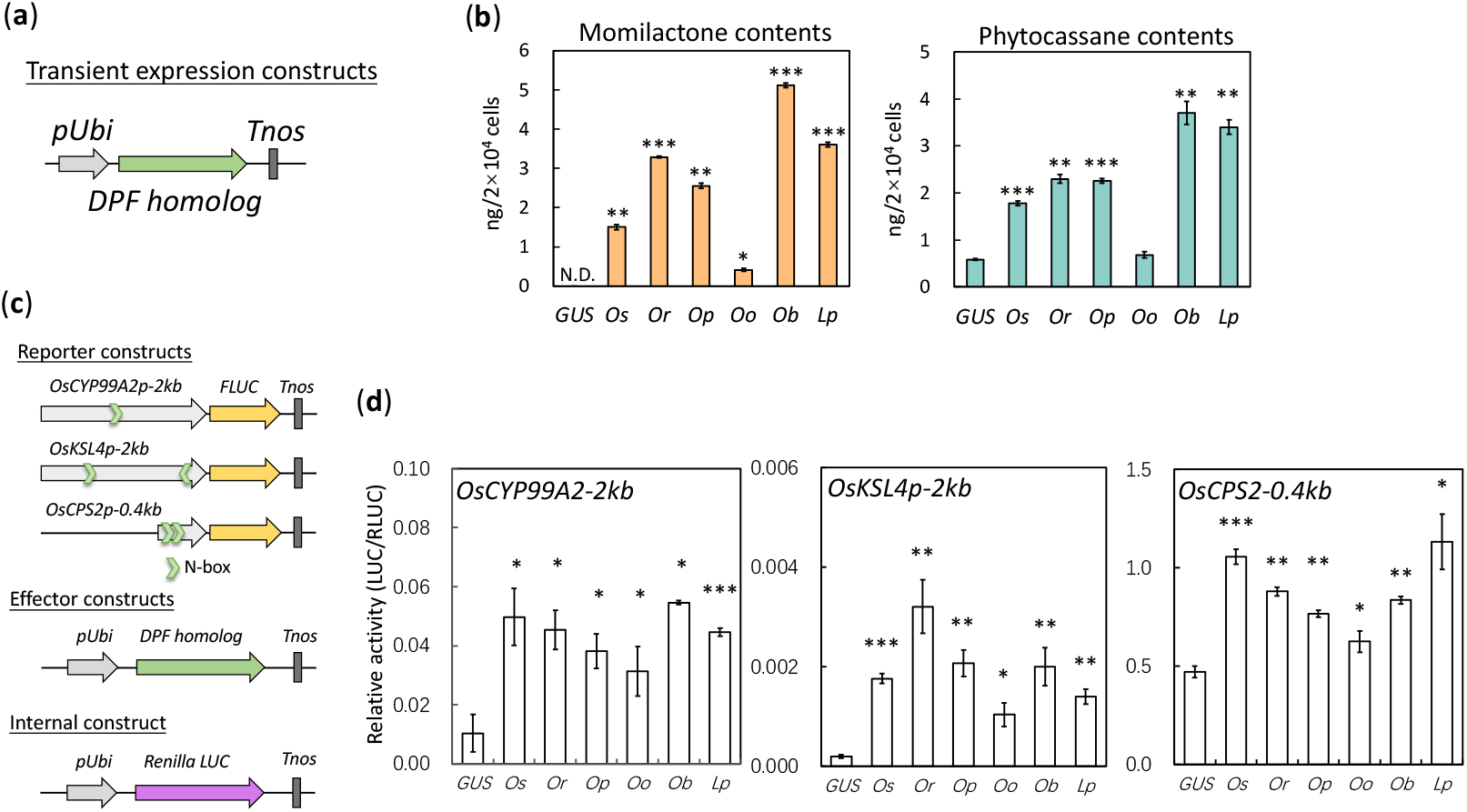
Transactivation activities of wild rice DPF homologues on diterpenoid phytoalexin production. (**a**) Schematic of transient expression vector. *DPF* homologue was driven by the maize ubiquitin promoter. (**b**) Accumulation of diterpenoid phytoalexins in *Oryza sativa* protoplasts expressing wild rice *DPF* homologues via transient expression vector. pUbi-*GUS*-Tnos was used as the negative control. Data represent the amount of momilactones or phytocassanes extracted from protoplasts (means ± SE, n = 3). *Os*, *Or*, *Op*, *Oo*, *Ob*, *Lp*: *DPF* and its homologues served as activators from *O. sativa*, *O. punctata*, *O. officinalis*, *O. brachyantha*, and *L. perrieri*. (**c**) Schematic of reporter vectors, effector vectors, and internal vector. (**d**) Results of reporter gene assay. Data indicate the relative luciferase (LUC) activities (firefly LUC/Renilla LUC) (means ± SE, n = 3). *Os*, *Or*, *Op*, *Oo*, *Ob*, *Lp*: *DPF* and its homologues served as effectors in the reporter assay. Asterisks indicate significant differences from the negative control (GUS) based on one-tailed Student’s t-test at *, *P* < 0.05, **, *P* < 0.01, ***, *P* < 0.001.

DPF presumably interacts with N-boxes in the promoter regions of *O. sativa* biosynthetic genes, including *OsKSL4*, *OsCPS2,* and *OsCYP99A2* (Yamamura *et al*., 2015). To investigate whether the wild rice DPF homologues can enhance the transcription of biosynthetic genes, we conducted a reporter gene assay using the DPF homologues as effectors. Three plasmids containing the promoter regions of *OsKSL4*, *OsCPS2*, and *OsCYP99A2* served as reporter vectors (Fig. 5c). The DPF homologues exhibited some variations in activity, but all of them significantly enhanced promoter activity in terms of the FLUC/RLUC ratio (Fig. 5d). The results at the transcription and metabolite levels strongly suggest the presence of a conserved *cis*-*trans* regulatory machinery for diterpenoid phytoalexin biosynthesis through DPF in *Oryza* and outgroup species.

### Analysis of genome-edited plant lacking N-boxes in the promoter region of *OsCPS2* and *OsKSL4* in *O. sativa*

The fact that DPF homologues are conserved in these wild rice species with functionality as trans-activators in diterpenoid phytoalexin biosynthesis in *O. sativa* inspired us to investigate the potential of *in planta cis*-acting sequences and how they conserve their promoter region in the genome, which is required for the *cis*-*trans* regulatory machinery of diterpenoid phytoalexin biosynthesis through DPF. Genome comparison of the upstream regions of diterpenoid phytoalexin biosynthetic genes revealed highly conserved regions between the cultivated and wild rice species. The comparison among *O. sativa*, *O. officinalis*, and *L. perrieri*, all of which are phytocassane producers, revealed a conserved common sequence for all three species at -400 bp to -500 bp upstream of the *O. sativa CPS2* start codon. These conserved regions in all three species contained an N-box *cis*-element (Fig. 6a). A similar analysis of the genome comparison in the upstream regions of *KSL4* in *O. sativa, O. punctata,* and *O. officinalis*, all of which are momilactone producers, showed an approximately 1 kb conserved region specifically between *O. sativa* and *O. officinalis*, which are split in two places on the corresponding upstream region of *O. officinalis KSL4* but located at almost one location with a small gap in the corresponding *O. sativa* genome (Fig. 6a). These results imply that it serves important functions as a *cis*-regulator region in the transcriptional regulation of the biosynthetic genes *KSL4* for momilactones and *CPS2* for phytocassanes.

**Fig. 6.**
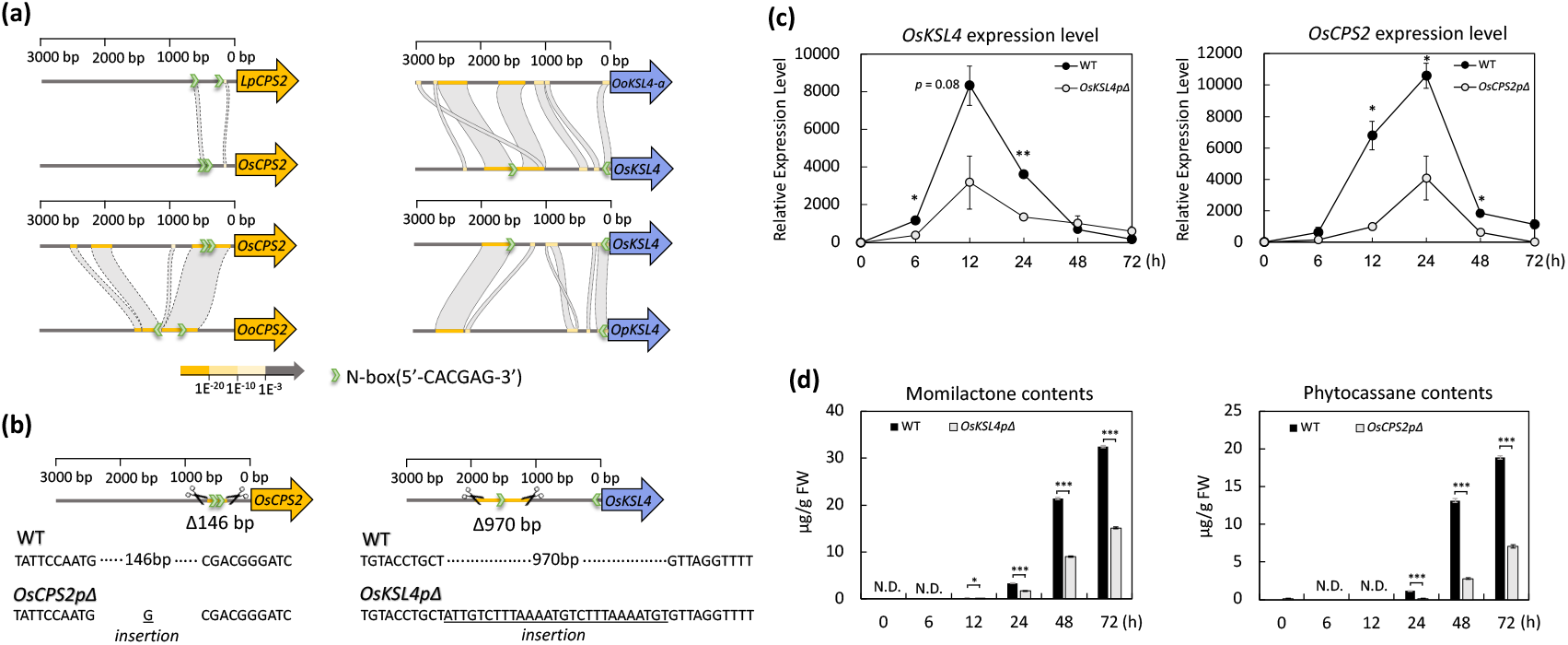
Effect of the deletion of N-boxes in the promoter regions of *OsKSL4* and *OsCPS2* on phytoalexin production. (**a**) 3 kb promoter regions of Os*CPS2* and Os*KSL4* homologues among cultivated and wild rice species. Conserved promoter fragments were colored in yellow color gradient. Green dots represent the N-box (5′-CACGAG-3′). (**b**) Promoter editing sites mediated by two individual gRNAs in two CRISPR-Cas9 editing vectors for each gene in cultivated rice (*Oryza sativa*). (**c**) Expression levels of *OsCPS2* and *OsKSL4* after time-course CuCl_2_ treatment in wild-type (WT) and genome-edited rice plant leaves. Left part: expression level of *OsKSL4* in WT and *OsKSL4pΔ* transgenic rice plants. Right part: expression level of *OsCPS2* in WT and *OsCPS2pΔ* transgenic rice plants. Data are shown as mean ± SE, n = 3. (**d**) Accumulation levels of momilactones and phytocassanes after CuCl_2_ time-course treatment. Black bars: Accumulation of momilactones or phytocassanes in WT. Gray bars: Accumulation of momilactones or phytocassanes in respective transgenic rice plants. Data are shown as mean ± SE, n = 3. Asterisks indicate significant differences in gene expression or phytoalexin production between the genome-edited and WT rice plants based on one-tailed Student’s t-test at *, *P* < 0.05, **, *P* < 0.01, ***, *P* < 0.001.

Notably, the N-box *cis*-element could only be found at close to the gap sequence at the border of the conserved region of the *KSL4* promoter in *O. sativa*. However, the upstream regions of *KSL4* in *O. officinalis* and *O. punctata* do not possess an exact N-box sequence. To investigate the detailed machinery between *trans*-activator DPF and *cis*-element N-box *in planta*, we generated two types of genome-edited *O. sativa* plants, which lack the conserved promoter regions containing the N-box *cis*-element on the promoter of *OsKSL4* (-1976∼-1006: Δ970bp, *OsKSL4pΔ*, with 27-base insertion) or *OsCPS2* (-482∼-336: Δ146bp, *OsCPS2pΔ*, with 1-base insertion) by using the CRISPR/Cas9 system. The *OsKSL4* promoter mutant was first selected as a heterozygous mutant in the T0 generation, and the homozygous mutant was successfully segregated in the T1 generation. For *OsCPS2* promoter mutant, we could obtain the homozygous mutant in the T1 generation from several lines of heterozygous mutant candidates (Fig. 6b). In addition, the MMCas9 cassette was lost from the genomes of these homozygous mutants; therefore, we used these MMCas9-free *OsKSL4* or *OsCPS2* promoter mutants in subsequent experiments (Fig. S1).

We treated genome-edited and WT rice leaves with CuCl_2_. The treatment significantly attenuated the expression levels of *OsKSL4* and *OsCPS2* in the *OsKSL4pΔ* and *OsCPS2pΔ* genome-edited rice plants compared with the WT plants (Fig. 6c). Correspondingly, the accumulation of momilactones and phytocassanes in the genome-edited transgenic rice plants decreased after CuCl_2_ treatment. These results suggest that loss of the N-box leads to the transcriptional attenuation of the biosynthetic genes of *OsKSL4* and *OsCPS2* under stress conditions. They also imply that other regulators are involved in the regulation of *OsKSL4* and *OsCPS2* because their expression could still be enhanced in genome-edited transgenic rice plants after CuCl_2_ induction. During flower development in these mutant plants, the accumulation level of momilactones significantly decreased in the *OsKSL4pΔ* mutant, whereas that of phytocassanes was not affected in the *OsCPS2pΔ* mutant compared with the WT plants (Fig. S2). These results suggest that the N-box in the promoter region of *OsKSL4* plays an important role in momilactone accumulation in the seeds, especially in the husks, during reproductive development, whereas the N-box in the promoter region of *OsCPS2* does not regulate phytocassane accumulation during seed development. These findings indicate that the N-box *cis*-sequence to which DPF binds has different functions in different tissues during rice diterpenoid phytoalexin synthesis under various environmental stress conditions.

## Discussion

In a previous study, we characterised the presence of conserved BGCs responsible for momilactone and phytocassane production in four wild rice species: *O. rufipogon*, *O. punctata*, *O. brachyantha*, *L. perrieri* (Miyamoto *et al*., 2016). However, the precise regulatory machinery governing diterpene biosynthesis in these wild rice species remains unclear. In the present study, we investigated the expression profile of biosynthetic genes, specifically *CPS* and *KSL* genes, after a time-course of treatment with CuCl_2_. Our findings revealed that this treatment induced the expression of biosynthetic genes and the accumulation of momilactone and phytocassane, as determined through LC-MS/MS analysis. These results strongly imply that these diverse wild rice species share a common regulatory framework for diterpene biosynthesis in response to abiotic stress.

*O. officinalis* is a CC genome-type wild rice species wherein we characterised momilactone and phytocassane BGCs in this study. Within the phytocassane BGC, we identified homologues of *CPS2*, *KSL7*, and several cytochrome *P450* genes in *O. officinalis*. Within the momilactone BGC, we observed a duplication of *KSL4* as well. After treatment with CuCl_2_, both *KSL4s* showed enhanced transcription. However, the precise roles of the duplicated *KSL4* genes in momilactone biosynthesis remain unclear. Furthermore, the momilactone levels after CuCl_2_ induction were significantly lower in *O. officinalis* than in the other rice species producing momilactones. Intriguingly, the expression level of *OoCPS4-c* was significantly induced after CuCl_2_ treatment. This result suggests that the diminished momilactone production originates from the insufficient transcriptional expression of other genes in the pathway (for instance, *OoKSL4s*), possibly implicating decreased metabolic flow, causing poor availability of precursor diterpene substrates.

In our RNA-Seq analysis of five distinct wild rice species after treatment with CuCl_2_, we systematically compiled stress-responsive transcription factors to elucidate the regulatory machinery governing diterpene biosynthesis. Among these transcription factors, the DPF homologues exhibited consistent transcriptional up-regulation in all five species. Notably, in *O. sativa*, DPF has been recognised as the dominant *trans*-activator in diterpenoid phytoalexin biosynthesis. This finding strongly suggests that DPF homologues play a pivotal role in regulating diterpenoid phytoalexin biosynthesis in wild rice species. To test this hypothesis, we first cloned the genes encoding DPF homologues into transient expression vectors, followed by their introduction into protoplasts, to assess diterpene production via LC-MS/MS analysis. The results provided conclusive evidence that all these homologues enhanced the production of diterpenoid phytoalexins. Furthermore, we confirmed their ability to activate the transcription of biosynthetic genes, such as *OsCPS2*, *OsKSL4*, and *OsCYP99A2*, through a reporter gene assay. These findings emphasise the roles of DPF and its homologues as universal *trans*-activators not only across the *Oryza* genus but also in related genera.

Protoplasts represent a potent tool for revealing physiological and molecular biological mechanisms in plants. Their versatile applications include the investigation of subcellular localisation and reporter gene assays (Zhang *et al*., 2011). A novel and expedient application of the protoplast platform involves the analysis of metabolites, facilitating a rapid assessment of the functions of transcription factors or enzymes associated with diterpenoid phytoalexin biosynthesis. In our efforts to identify wild rice homologues of *O. sativa* transcription factors associated with diterpenoid biosynthesis, we found the homologues *OsTGAP1*, *OsbZIP79*, and *OsWRKYs*, which were conserved across wild rice species but not included in our DEG list (Fig. S3). In fact, they did not exhibit significant responsiveness to CuCl_2_ treatment in foliar parts as the *DPF* homologues. Two bZIP factor genes, *OsTGAP1* and *OsbZIP79*, were mainly expressed in the roots; thus, they were not identified as DEGs in our transcriptome analysis of foliar pieces. Of the transcription factors that were strongly correlated with the accumulation of diterpenoid phytoalexins, only DPF and OsTGAP1 enhanced the production of diterpenoid phytoalexins by a single introduction into protoplasts (Fig. S4). Both of these transcription factors induced phytoalexins in stable overexpression plants, even in the absence of stress, suggesting that using a protoplast assay system is a concrete method to screen for phytoalexin inducers in a short period of time (Yamamura *et al*., 2015; Okada *et al*., 2009).

Notably, *O. brachyantha* is uncapable of momilactone and phytocassane biosynthesis because of incomplete BGCs. Nevertheless, the DPF in *O. brachyantha* retained its *trans*-activation activity in diterpenoid phytoalexin biosynthesis in *O. sativa* protoplasts, making the precise function of *ObDPF* in *O. bracyantha* an interesting subject for future investigation. In addition, the gene cluster loci in *O. brachynatha* chromosomes share the smallest cluster size (approximately 100 kb), containing two to four P450 gene homologues of *OsCYP76M* and *OsCYP99A* in the clusters, which are highly induced by CuCl_2_ treatment. Whether *ObDPF* is involved in the regulation of such *P450* genes in the clusters remains to be clarified, but the existence of N-boxes in the promoter regions of these *P450* genes in the cluster implies that they are likely to be transcriptionally regulated by *ObDPF* (Fig. S5). In our future work, we will conduct a transactivation analysis of *ObDPF* in the promoter regions lacking N-boxes in the *O. bracyhantha* genome to elucidate the molecular mechanisms underlying the transcriptional regulation of the cluster region. Moreover, DPF homologues not only exist in *Oryza* and closely related genera, such as *Leersia*, but have also been found in wheat (*Triticum aestivum*) and even in species from the PACMAD clade, implying that DPF homologues regulate as-yet-unknown diterpenoid phytoalexin biosynthesis in Poaceae (Fig. S6).

The N-box has been identified as the presumed binding motif for DPF (Yamamura *et al*., 2015). We investigated the N-box distribution in the 3 kb promoter regions of biosynthetic genes in wild and cultivated rice species. The results showed that the N-boxes in the cluster regions were slightly more enriched than those in the whole genome regions (e.g., in the case of *O. sativa*, 61.3% of the genes within clusters shared N-boxes in the promoter, whereas 58.3% of the genes in the whole genome had N-boxes). However, the results of Fisher’s exact test did not show a high frequency (Dataset S6). As shown in Fig. S5, not all but many of the clustered genes in either cultivated or wild rice have N-boxes and G-boxes (5 ′-CACGTG-3 ′), which is a typical *cis*-acting element for a type of bHLH factor, indicating that N-boxes and related *cis*-acting elements are recruited to regulate diterpenoid phytoalexin biosynthetic genes in the clusters (Fig. S5). An exception was the absence of N-boxes in the conserved upstream region of the *KSL4* homologue in *O. puncutata* and *O. officinalis*, suggesting that a binding motif of a DPF homologue other than N-boxes regulates momilactone production in these wild rice plants. In fact, in the upstream region of *OoKSL4* has a similar sequence (5′-T<u>AC</u>A<u>AG</u>-3′) to the N-box (5′-C<u>AC</u>G<u>AG</u>-3′), which is surrounded by a highly conserved sequence to the *OsKSL4* promoter (Fig. S7). Regarding the CPS2 promoter, the upstream region of *OsCPS2* has a tandem arrangement of two N-boxes, but one in the corresponding *OoCPS2* promoter region reveals a single nucleotide substitution (5′-<u>CA</u>T<u>GAG</u>-3′) to the N-box. These observations suggest that *O. officinalis* has some diversification of N-box-related sequences in such highly conserved upstream regions of diterpenoid phytoalexin biosynthetic genes, such as *KSL4* and *CPS2*. This fracturation of *cis*-sequences would explain why *O. officinalis* produces a low amount of diterpenoid phytoalexins. Although we cannot draw a solid conclusion on the evolutionary trajectory of the *cis*-sequence, the evolutionary nucleotide transitions in this region possibly affect gene expression and consequently alter diterpenoid phytoalexin production. Another possibility is that DPF does not act directly as a *trans*-activator in wild rice to control *KSL4* expression but is involved in the regulation of this process via other transcription factors mediated by DPF.

In this study, we generated genome-edited rice plants using *O. sativa*, resulting in rice a plant with a 970 bp deletion (but 27 bp insertion) in the *OsKSL4* promoter and a rice plant with a 146 bp deletion (but 1 bp insertion) in the *OsCPS2* promoter. The insertion of a small fragment was also observed when CRISPR-Cas9 was used to delete genome fragments (Wang *et al*., 2017; Cai *et al*., 2018). The expression levels of both biosynthetic genes were significantly down-regulated in the genome-edited rice plants but were not completely abolished after CuCl_2_ treatment. This result implies that the N-box motif and DPF *cis*-*trans* interaction are important for the regulation of the defense response, but they do not seem to be the only factors involved in the inductive regulatory mechanism of diterpenoid phytoalexin accumulation.

Interestingly, *L. perrieri* exhibited super-overrepresented N-boxes in the phytocassane gene cluster regions on chromosome 2 (81.8% of the genes on the cluster versus 61.1% of the genes on the whole genome have N-boxes in their promoter regions), whereas it did not have either genes or N-boxes in the region corresponding to the momilactone BGC on chromosome 4 in *Oryza* species (Fig. S5). This example also fits the situation in which the frequency of N-box occurrence correlates well with the production level of phytocassanes, which is almost in the same range as that in *O. sativa* and *O. rufipogon.* Transformation of wild rice, including genome editing, is currently under investigation, and the importance of regulatory regions and *cis* sequences in wild rice as a host is expected to be clarified in the future.

The integration of genomic and transcriptomic analyses of wild crops has significantly contributed to deciphering the regulatory mechanisms of diterpenoid and phytoalexin biosynthesis in rice plants. Furthermore, as delineated in this study, a firm examination of the contribution of *cis* sequences and their relationship to *trans* factors is a definite step toward the full elucidation of the sophisticated regulatory mechanisms of diterpenoid phytoalexin production in rice plants. Our findings provide insights into the evolutionary footprint of the regulatory mechanisms of diterpenoid phytoalexins by *cis*-*trans* factors, which would have been built along the BGC evolutionary track from wild to cultivated rice.

## Acknowledgements

We are grateful to Dr. Matthew R Shenton (Institute of Crop Science, NARO) for providing the *O. officinalis* genome sequence data before its publication. We also thank Dr. Masaki Endo (Institute of Agrobiological Sciences, NARO, NARO) for providing the genome-editing vectors pU6gRNA and pZH_gYSA_MMCas9. This work was supported in part by JSPS KAKENHI grant no. 20H02922, 24K01690, Japan-Austria Research Cooperative Program between JSPS and FWF (grant number JPJSBP120202002), NIG-JOINT (15A2021) to KO and a project of the Science and Technology Department of Sichuan Province (2020-YFH0161), a project of the Science and Technology Department of Chengdu City (2021-GH02-00085-HZ) to NY. The wild rice accessions used in this study were distributed from the National Institute of Genetics supported by the National Bioresource Project, MEXT, Japan. We would like to thank Editage (www.editage.jp) for English language editing.

## Competing interests

The authors declare no conflicts of interest directly relevant to the content of this article.

## Author contributions

KO and HN conceived the study. YS and HF prepared the wild rice materials. YT and MK performed RNA sequencing. NY and AZ conducted transcriptomic data analysis. YL, IM, ST, KM, and MM performed molecular cloning, genome editing, and functional analyses. KO and HN supervised this study. YL drafted the manuscript, and KO revised the manuscript with contributions from all authors. All the authors have read and approved the final version of the manuscript.

## Supporting Information

**Fig. S1.**
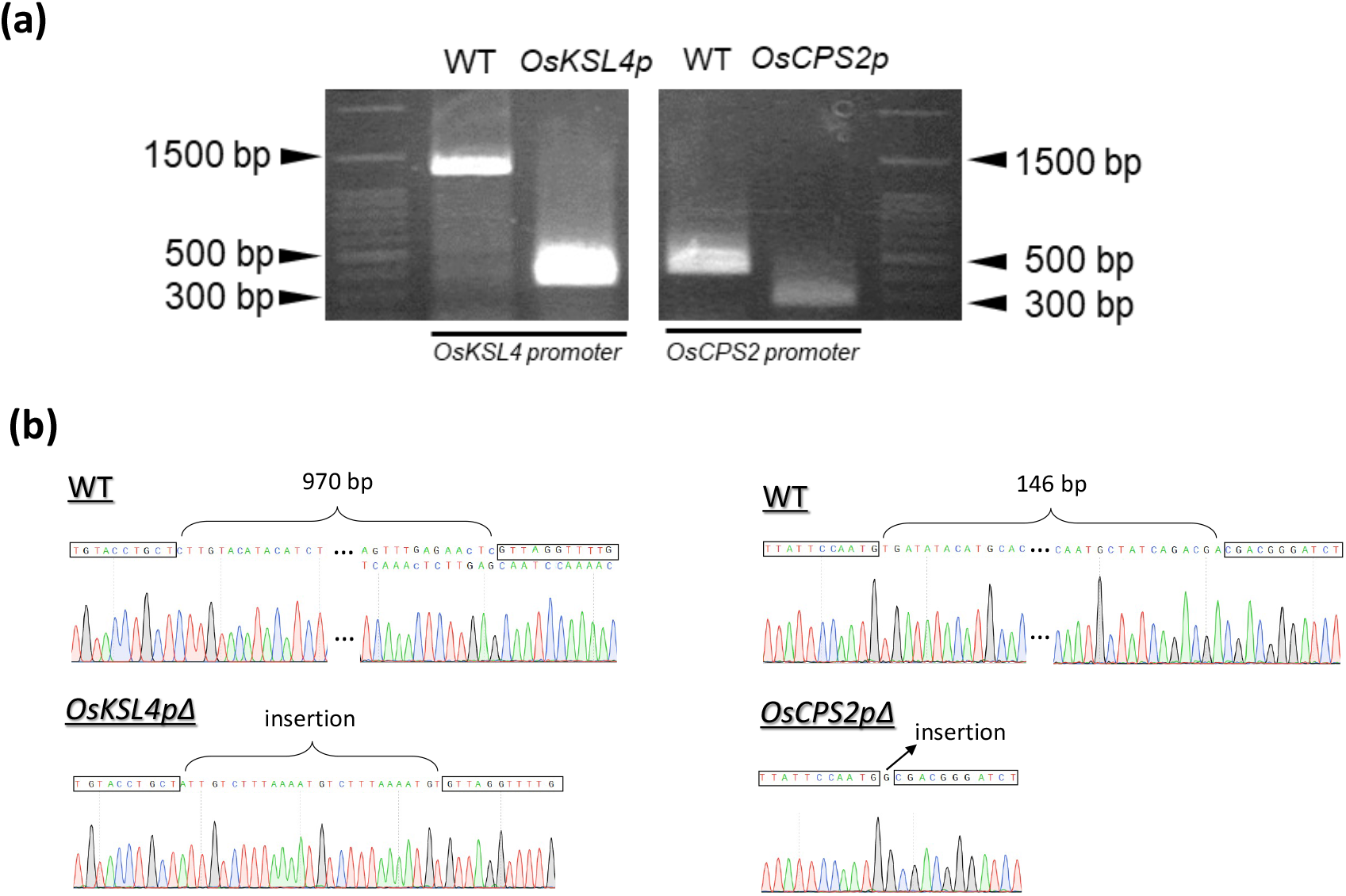
Gel image and sequencing chromatograms of *OsCPS2* and *OsKSL4* promoter amplification by polymerase chain reaction (PCR) from DNA samples of wild-type (WT) and genome-edited rice plants. (a) Expected intact fragment sizes are 1298 bp for *OsCPS2* and 428 bp for *OsKSL4* in WT rice. The expected mutated fragment size is 328 bp for *OsKSL4* in *OsKSL4p* genome-edited transgenic rice plants and 282 bp for *OsCPS2* in *OsCPS2p* genome-edited transgenic rice plants. (**b**) Sequencing chromatograms of WT and genome-edited rice plants.

**Fig. S2.**
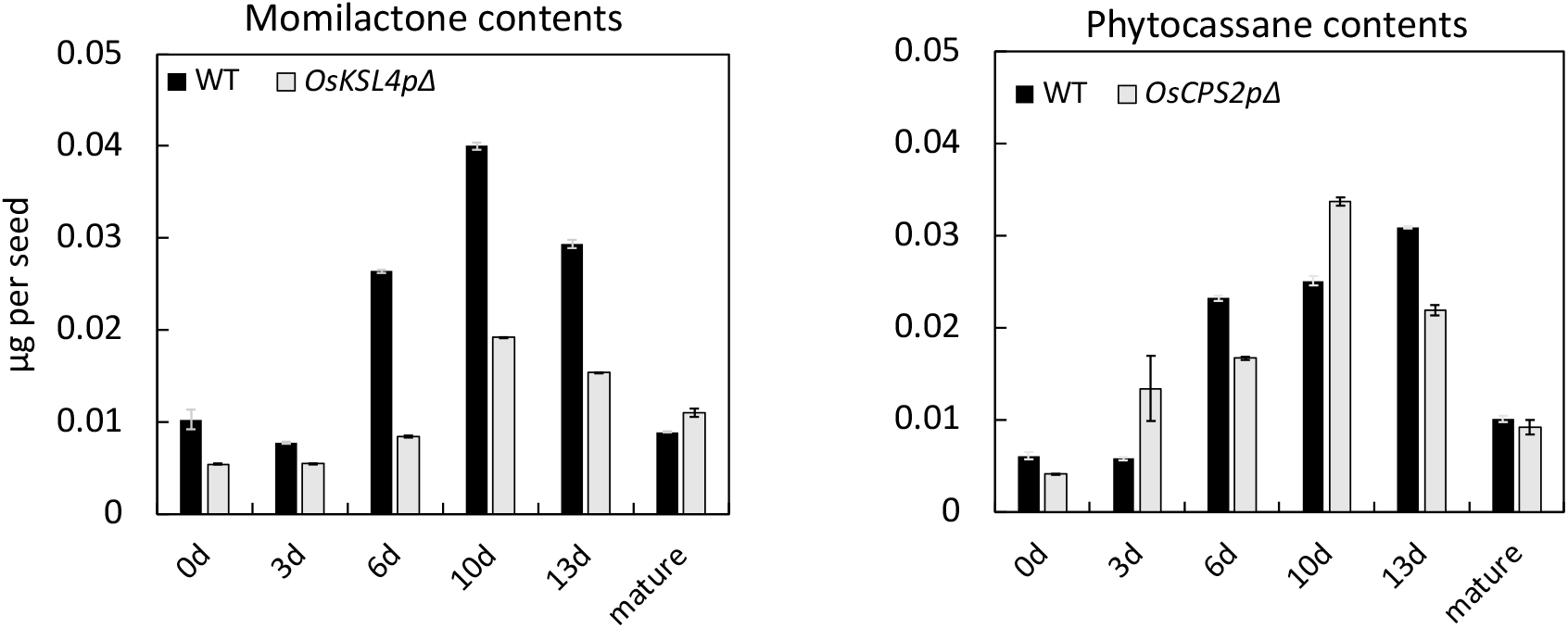
Accumulation level of momilactones and phytocassanes in seeds at the ripening stage. Black bars: Accumulation of momilactones or phytocassanes in WT. Gray bars: Accumulation of momilactones or phytocassanes in respective transgenic rice plants. Data are shown as mean ± SE, n = 3.

**Fig. S3.**
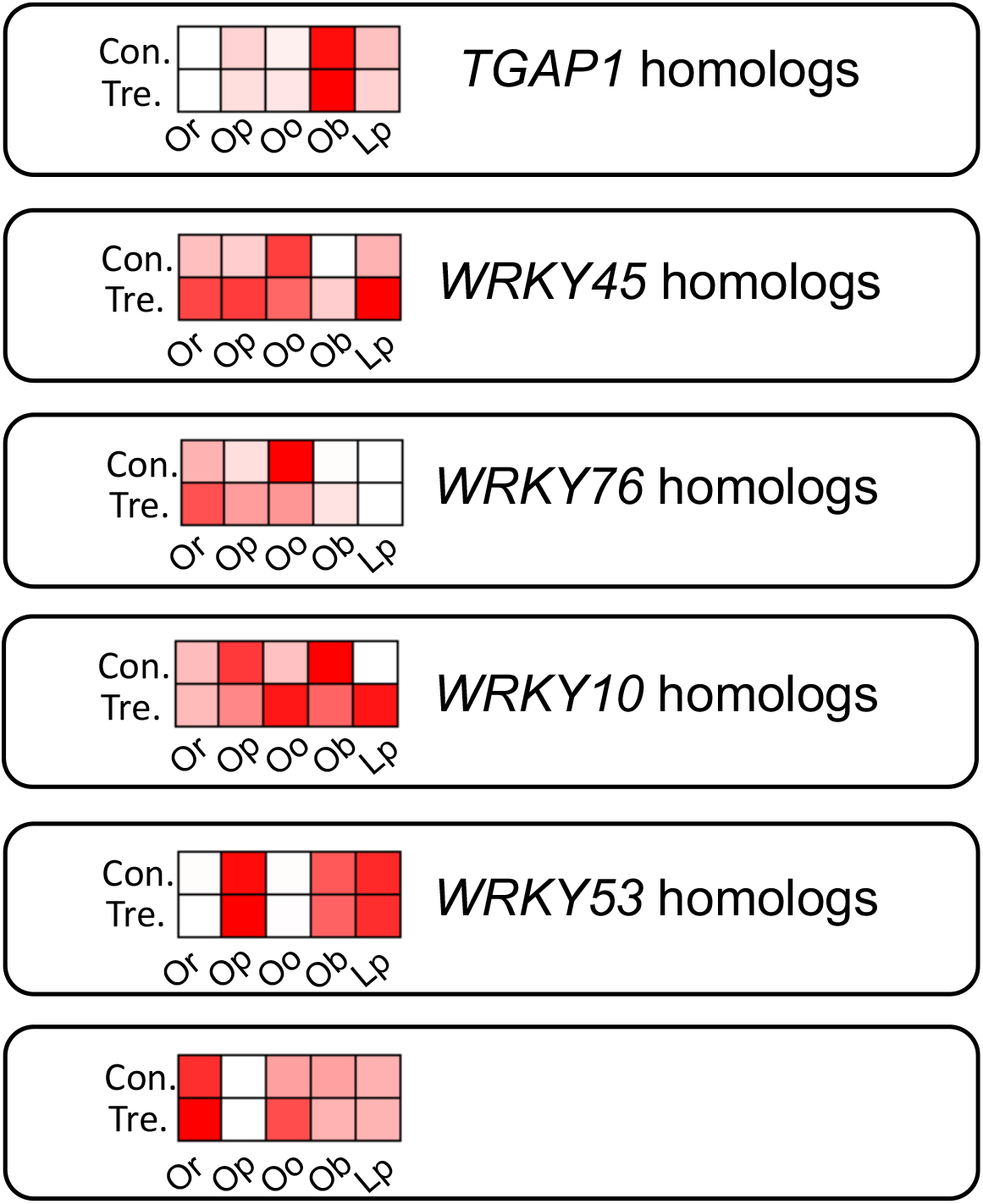
Expression of homologues of *Oryza sativa* transcription factors related to diterpenoid phytoalexin biosynthesis. Protein sequences of *O. sativa* transcription factors served as query and searched in protein sequence data in each wild rice species. Heatmap represents expression profiles in the control and treatment groups after 24 h of CuCl_2_ treatment. Transcripts per million (TPM) (log_2_ scale) is shown in the heatmap. Red boxes represent high expression levels, while white boxes represent low expression levels. Con.: control group, Tre.: treatment group. *Or*: *Oryza rufipogon*, *Op*: *Oryza punctata*, *Oo*: *Oryza officinalis*, *Ob*: *Oryza brachyantha*, *Lp*: *Leersia perrieri*.

**Fig. S4.**
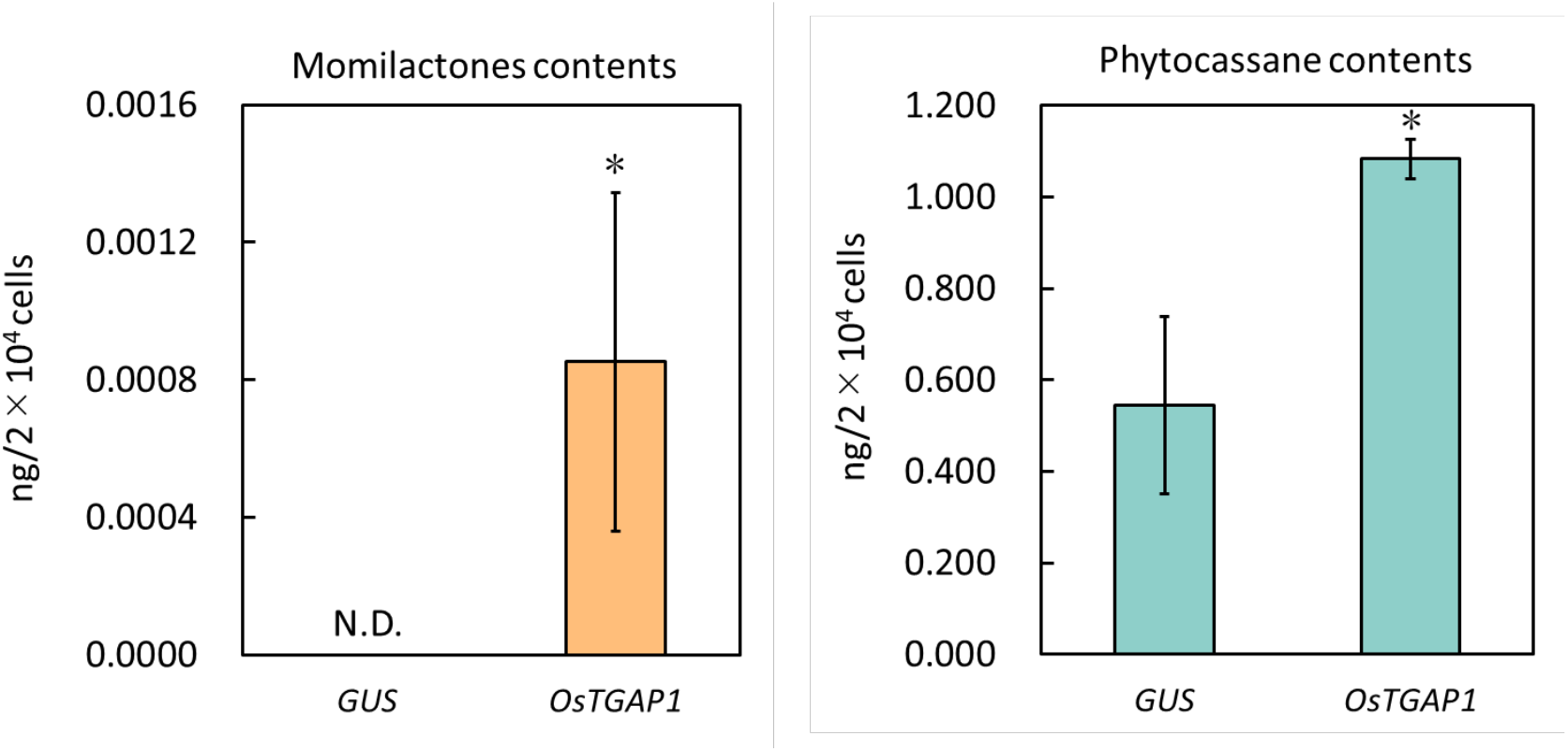
Accumulation of diterpenoid phytoalexins in *Oryza sativa* protoplasts expressing *OsTGAP1* via transient expression vector. pUbi-*GUS*-Tnos was used as a negative control. Data represent the amount of momilactones or phytocassanes extracted from protoplasts (means ± SE, n = 3). Asterisks indicate significant differences from the negative control (*GUS*) based on one-tailed Student’s t-test at *, P < 0.05, **, P < 0.01, ***, P < 0.001.

**Fig. S5.**
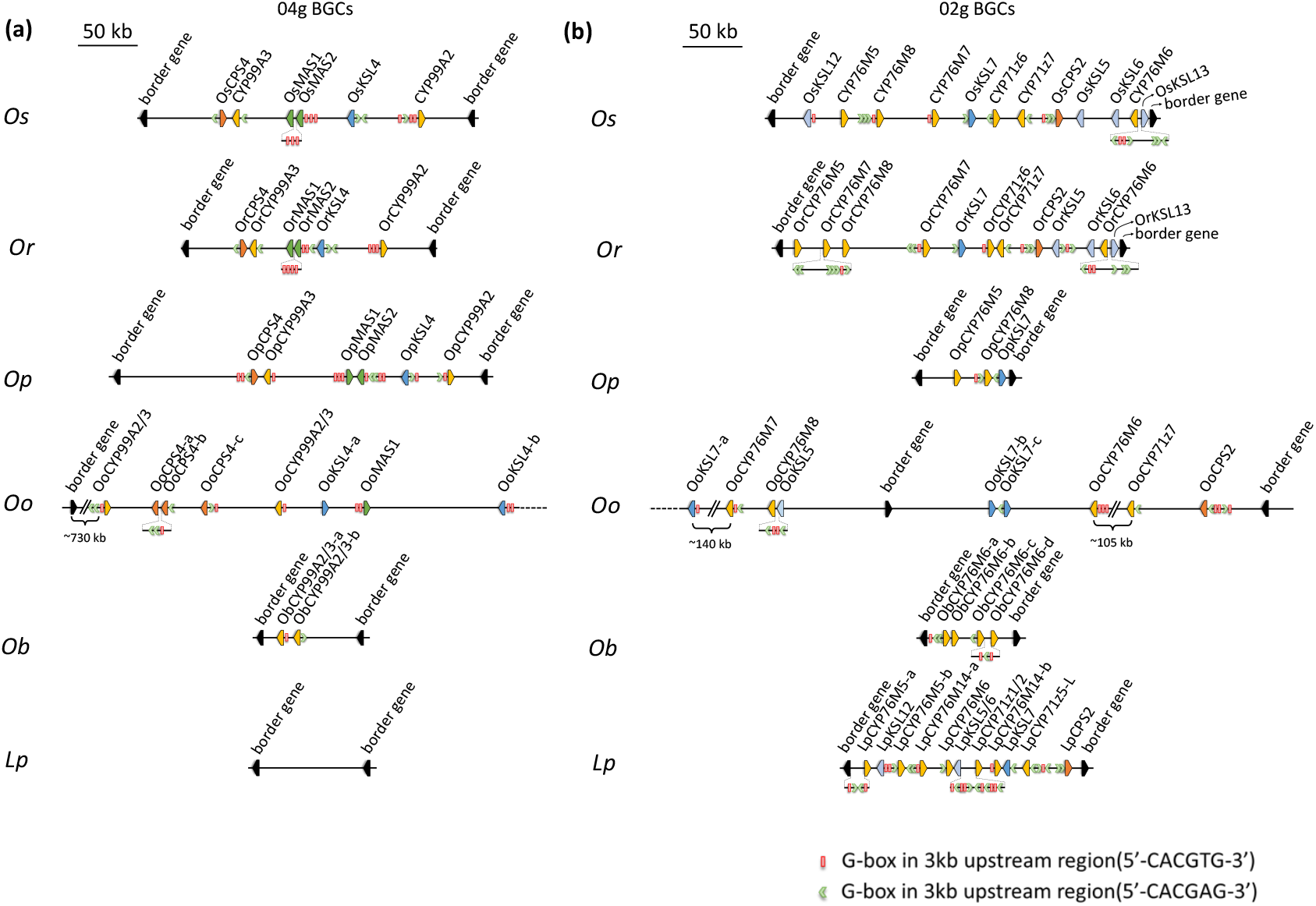
Schematic of gene loci in cultivated rice *Oryza sativa* and wild rice biosynthetic gene clusters (BGCs) on correspondent chromosomes 2 and 4 with N-boxes in their upstream regions. (a) momlacotone biosynthetic cluster on chromosome 4. (b) phytocassane biosynthetic cluster on chromosome 2. *Os*: *Oryza sativa*, *Op*: *Oryza punctata*, *Oo*: *Oryza officinalis*, *Ob*: *Oryza branchyantha*, *Lp*: *Leersia perrieri*. Green arrows represent the location of the N-box (5′-CACGAG-3′) in the upstream regions of the biosynthetic genes. Red blocks represent the location of palindrome *cis*-element G-box (5′-CACGTG-3′).

**Fig. S6.**
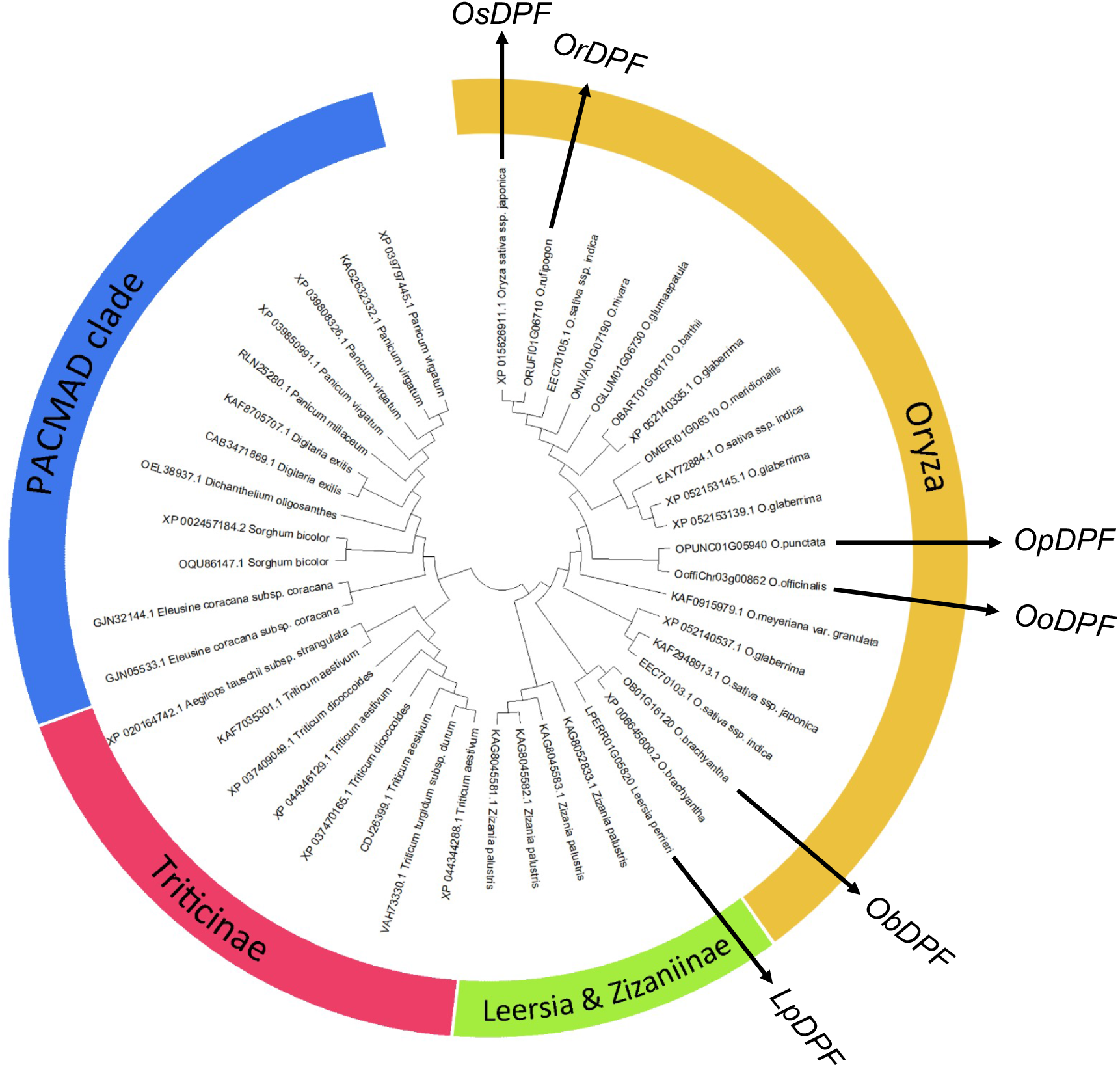
Phylogenetic relationship of diterpenoid phytoalexin factor (DPF) homolog proteins from Poaceae. Phenogram representation of DPF homolog proteins from Poaceae using the Maxima Parsimonia method. Multiple sequence alignment was generated using the MUSCLE algorithm, and a phylogenetic tree was constructed using the MEGA X program (Kumar et al., 2018). Proteins from the *Oryza* genus are indicated in yellow, proteins from *Leersia* and *Zizaniinae* in green, proteins from *Triticinae* in red, and proteins from the PACMAD clade in blue.

**Fig. S7.**
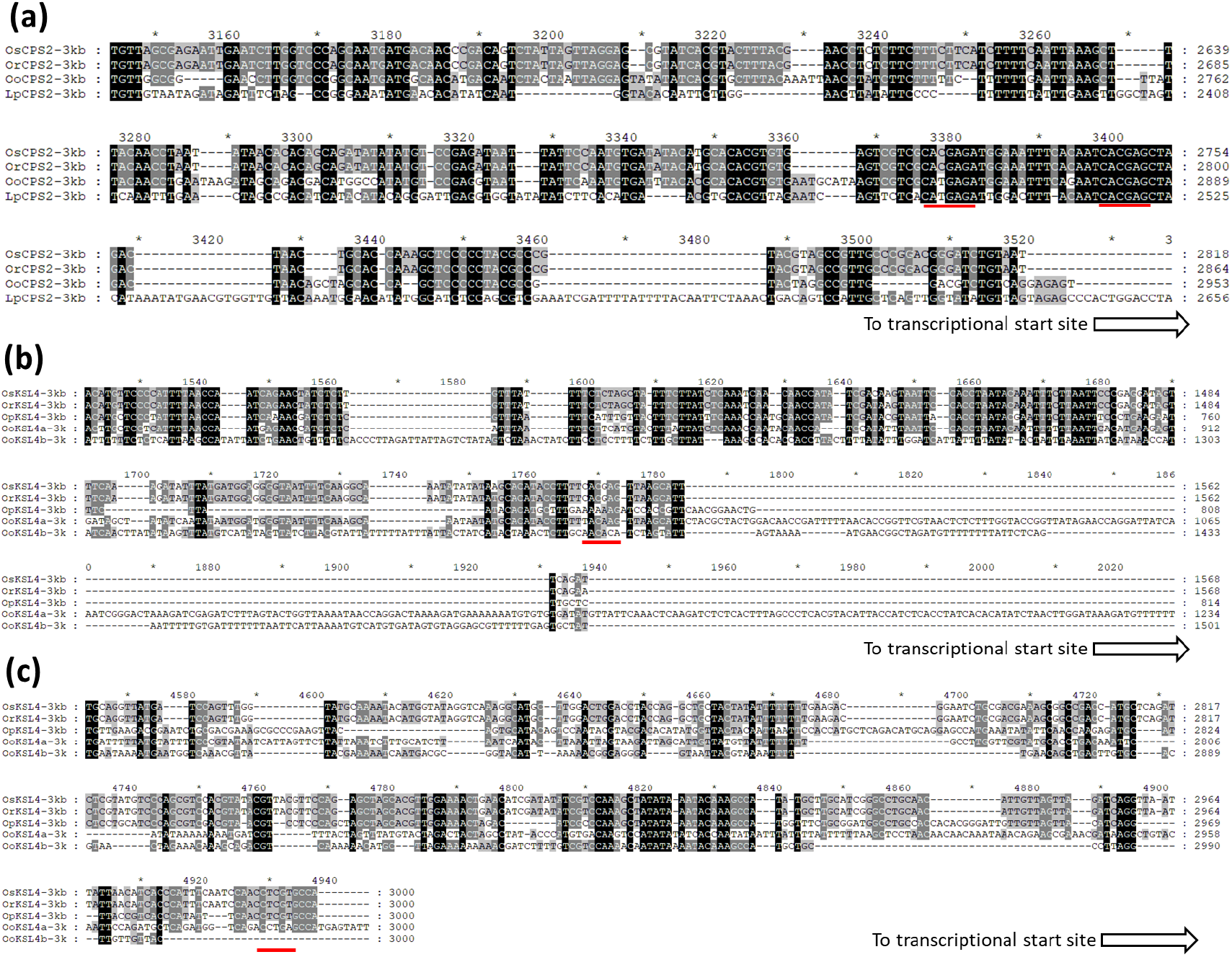
Alignment of upstream sequences of CPS2 and KSL4 from *Oryza sativa* and wild rice, which possess highly conserved regions with N-boxes. Alignment results of the *CPS2* promoter region containing N-boxes (5′-CACGAG-3′). (b) Alignment result of the *KSL4* promoter region containing N-boxes. (c) Alignment result of the *KSL4* promoter region containing N-boxes on reverse strain. *Os*: *Oryza sativa*, *Or*: *Oryza rufipogon*, *Op*: *Oryza punctata*, *Oo*: *Oryza officinalis*, *Lp*: *Leersia perrieri*. Red lines indicate the location of N-boxes.

**Table S1.** Gene locus and symbols in *Oryza officinalis*.

| Chr02. |  |
| --- | --- |
| *OoffiChr02g01875* | *OoKSL7-a* |
| *OoffiChr02g01876* | *OoCYP76M7* |
| *OoffiChr02g01877* | *OoCYP76M8* |
| *OoffiChr02g01878* | *OoKSL5* |
| *OoffiChr02g01880* | *border gene* |
| *OoffiChr02g01883* | *OoKSL7-b* |
| *OoffiChr02g01884* | *OoKSL7-c* |
| Manually annotated gene | *OoCYP76M6* (Chr04:38909020-38910525) |
| *OoffiChr02g01887* | *OoCYP71z7* |
| *OoffiChr02g01888* | *OoCPS2* |
| *OoffiChr02g01891* | *border gene* |
| Chr04. |  |
| *OoffiChr04g00260* | *OoCYP99A2/3* |
| *OoffiChr04g00261* | *OoCPS4-a* |
| *OoffiChr04g00262* | *OoCPS4-b* |
| *OoffiChr04g00263* | *OoCPS4-c* |
| *OoffiChr04g00264* | *OoCYP99A2/3* |
| *OoffiChr04g00265* | *OoKSL4-a* |
| *OoffiChr04g00266* | *OoMAS-1* |
| *OoffiChr04g00269* | *OoKSL4-b* |

**Table S2.**
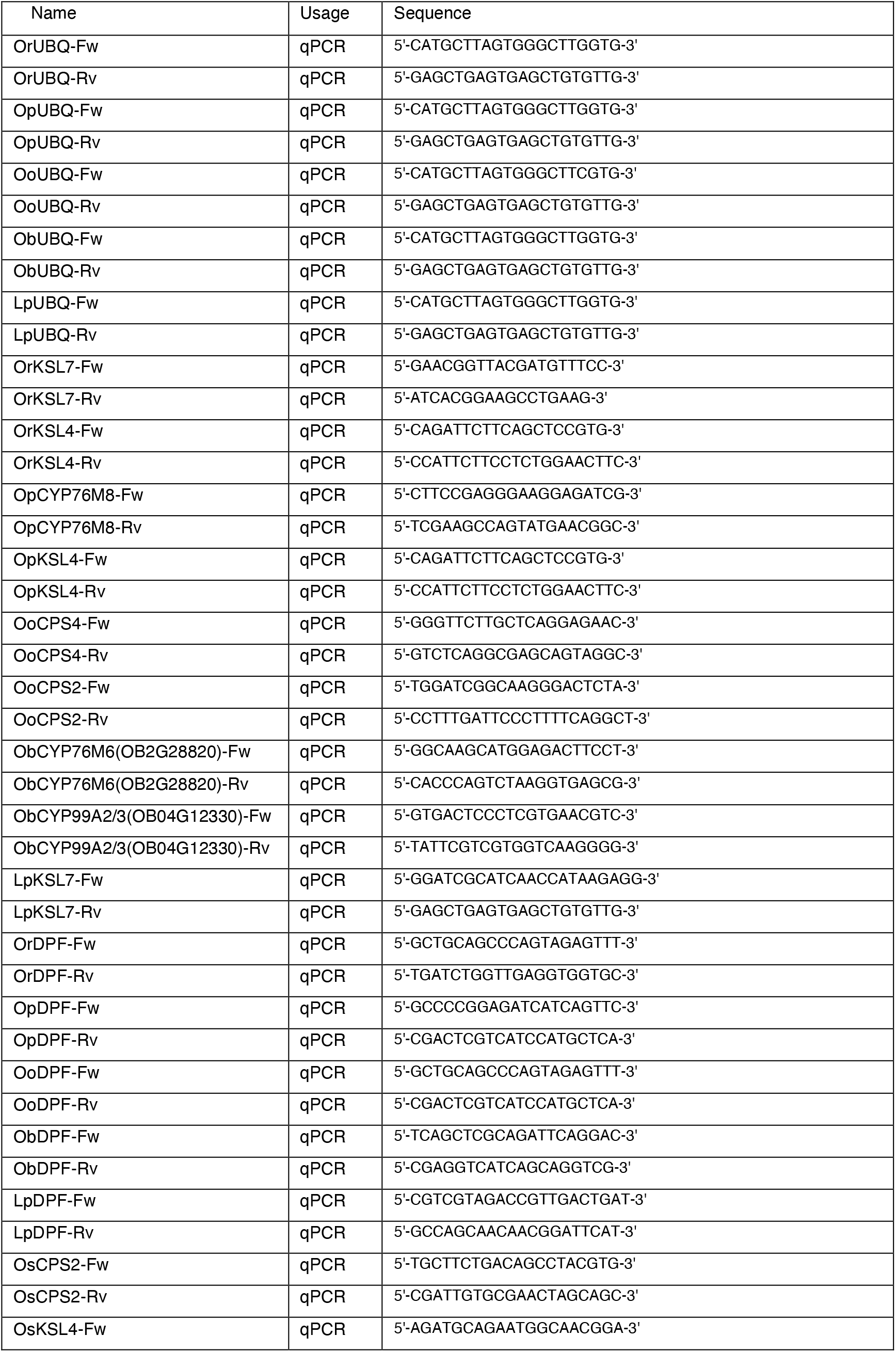
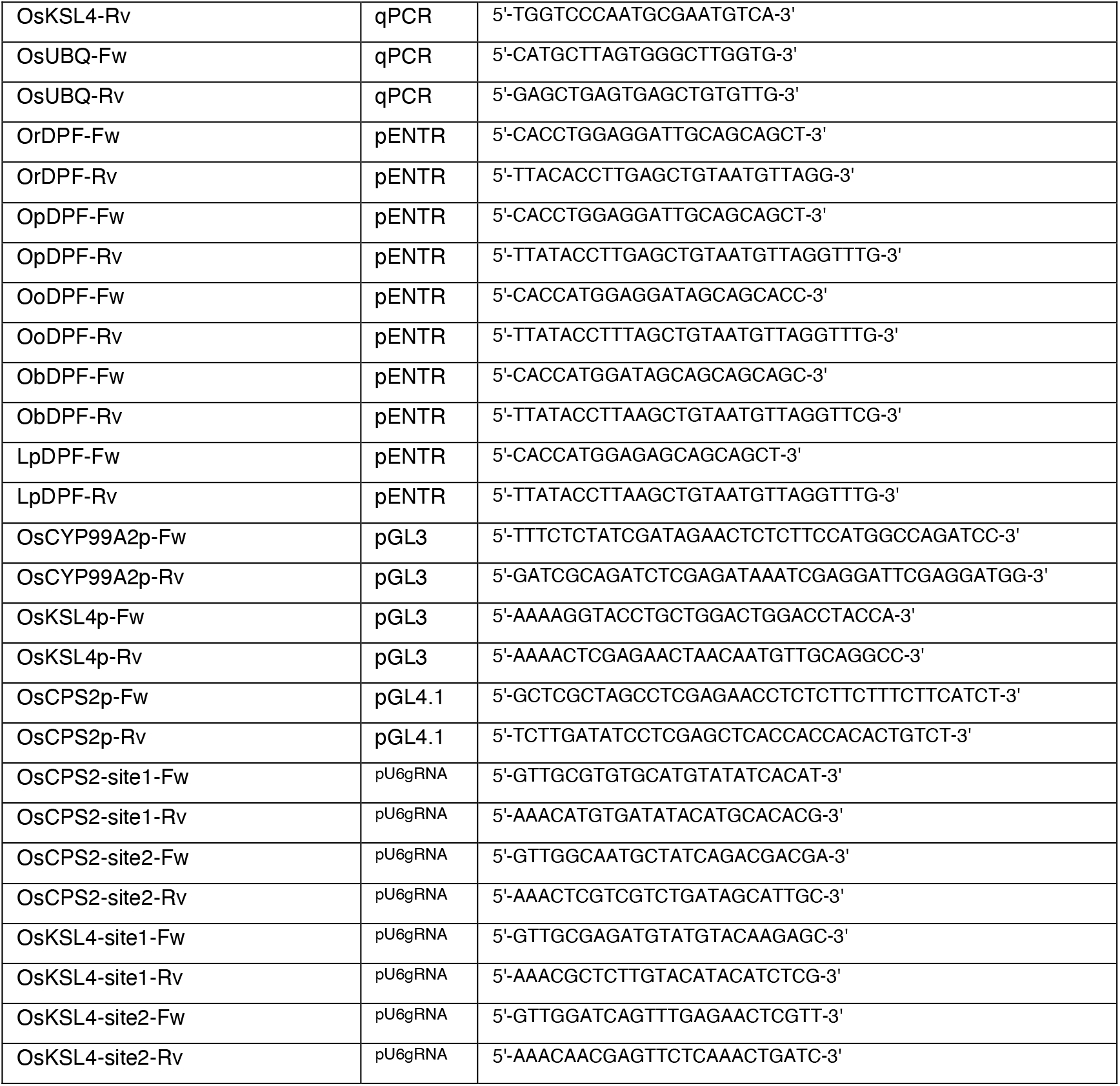
Primers used in this study.

**Dataset S1** Up-regulated DEGs among five wild rice species.

**Dataset S2** Overrepresented GO terms in up-regulated DEGs after CuCl_2_ treatment.

**Dataset S3** Over-represented gene modules in the up-regulated DEGs after CuCl_2_ treatment.

**Dataset S4** Up-regulated transcription factors after CuCl_2_ induction.

**Dataset S5** Coding sequences of diterpenoid phytoalexin factor (DPF) homologues among five species.

**Dataset S6** Chi-square test of N-boxes and G-boxes in 3 kb upstream regions in wild and cultivated rice.

## Abbreviation

BGC: Biosynthetic gene cluster
bHLH: Basic helix-loop-helix
CuCl_2_: Copper chloride
DEGs: Differentially expressed genes
DPF: Diterpenoid phytoalexin factor
FDR: False discovery rate
GGDP: Geranylgeranyl diphosphate
UV: Ultraviolet
*Or, O. rufipogon*: *Oryza rufipogon*
*Op, O. punctata*: *Oryza punctata*
*Oo, O. officinalis*: *Oryza officinalis*
*Ob, O. branchyantha*: *Oryza brachyantha*
*Lp, L. perrieri*: *Leersia perrieri*
GO: Gene Ontology
*GUS*: *β-glucuronidase*
RT-qPCR: Reverse transcription-quantitative polymerase chain reaction
*RLUC*: *Renilla luciferase*
TPM: transcripts per million
*LUC, FLUC*: *Firefly luciferase*
LC-MS/MS: Liquid chromatography with tandem mass spectrometry
WT: Wild type

## Notes

### Competing Interest Statement

The authors have declared no competing interest.

