## Supplementary Dataset 5 for "Evolutionary conserved *cis-trans* regulation machinery for diterpenoid phytoalexin production in Poaceae"

>OsDPF

ATGGAGGATTGCAGCAGCTGGATCCATGGCTACGCCAACGCTAACGCCACCGCCGGCAACAACGGCTTCATGTGCGGCTACGCTGCCAGCTGCAGCCCAGTAGAGTTTCAGCAGCAGCAACAGCTGGTCGGCTCGCAGATTGAGCACCACCTCAACCAGATCAGCATGCAGATGGGGATGGATGACGAGTCGGCGGTGTACGACGGCGCCTCCATGGTGGACGTCCTCCTCATGGCTTCCTCGTCGCCGCACCACCACGCCGGCGCCGGCAGCTTCCAGTACTCCTCGCCGACGTCCTCCTCCGCCTCCTTCCGCTCCGCCTCCGTCTCGTGCAGCCCTGAGAGCTCGGCGGCGGCGACGACGCACTTCCTCGGACCGCCGGCGCCGTCCGCGGCGGCGGCGGGGTTCCATTACCCGGAGGTCTCCTCGCAGGCGCCGTTGCCACTACCCTTGCCGCCCTACGAGCCGCAGCACGGCCAATACACCACCGTCCTCTCGCCGCCGCCGCCGGCGCCAGAGTTGCCGGCGACTACTACGCCGGCGACCGGCGGCGCGTTCAGGCGGTACGCGCGGCACCTCCGCCCGAGGAGGCTGCCCAAGCCGGGAGGGTGCGGGCAGAGGATGTTCAAGACGGCCATGTCGGTGCTCACCAAGATGCACGTGGCGGCGACGTACAACCGCCAGTACTACTACCAGCAGGCGGCAGCCGCCGCCGCGTCGGCGTCGGCGGCCGAGGCGCCGCCGTCCGGCAACCAGCTGCAGCACATGATCTCGGAGCGGAAGCGGCGGGAGAAGCTCAACGACAGCTTCCTCGCCCTCAAGGCCGTCCTCCCTCCCGGCTCTAAGAAAGACAAGACATCGATACTGATCAGAGCAAGAGAGTACGTAAAATCTCTCGAGTCAAAGCTGTCAGAGCTGGAGGAGAAGAACCGGGAGCTAGAGGCGAGGCTAGCCAGCCGCCCCGCCGCCGCCGCCAAGAACGACAAAGGCGAGACGGCGGCGGCGCCAGCGCCGGAAGCCGGCGATGAGACGAAGCGAAAGGACCTAGTAGAGATCGAGGTGACGACGAGCGGCGGCGGCGCCGGAGCGGCGGATGCGGCGGCGGCGGCGGGAGGAGATCAAGAGACCTGCACGCTCAACGTAGACTTGCGCGGCGGCGGAGGCGGCGGAGGCATGAGCACGACGGACGTTGTGCTCCGGACGCTGCAGTGCCTGAGAGAGCAGATCGGCGACGGCGCCAGCCTCGTGGCGATGAGCACCAGCGCCGGCTCCGGCGGCCGCCCGCCTCGTGCAAACCTAACATTACAGCTCAAGGTGTAA

>OrDPF

ATGGAGGATTGCAGCAGCTGGATCCATGGCTACGCCAACGCTAACGCCACCGCCGGCAACAACGGCTTCATGTGCGGCTACGCTGCCAGCTGCAGCCCAGTAGAGTTTCAGCAGCAGCAACAGCTGGTCGGCTCGCAGATTGAGCACCACCTCAACCAGATCAGCATGCAGATGGGGATGGATGACGAGTCGGCGGTGTACGACGGCGCCTCCATGGTGGACGACCTCCTCATGGCTTCCTCGTCGCCGCACCACCACGCCGGCGCCGGCAGCTTCCAGTACTCCTCGCCGACGTCCTCCTCCGCCTCCTTCCGCTCCGCCTCCGTCTCGTGCAGCCCTGAGAGCTCGGCGGCGGCGACGACGCACTTCCTCGGACCGCCGGCGCCGTCCGCGGCGGCGGCGGGGTTCCATTACCCGGAGGTCTCCTCGCAGGCGCCGTTGCCACTACCCTTGCCGCCCTACGAGCCGCAGCACGGCCAATACACCACCGTCCTCTCGCCGCCGCCGCCGGCGCCAGAGTTGCCGGCGACTACTACGCCGGCGACCGGCGGCGCGTTCAGGCGGTACGCGCGGCACCTCCGCCCGAGGAGACTGCCCAAGCCGGGAGGGTGCGGGCAGAGGATGTTCAAGACGGCCATGTCGGTGCTCACCAAGATGCACGTGGCGGCGACGTACAACCGCCAGTACTACTACCAGCAGGCGGCGGCCGCCGCCGAGTCGGCGTCGGCGGCCGAGGCGCCGCCGTCCGGCAACCAGCTGCAGCACATGATCTCGGAGCGGAAGCGGCGGGAGAAGCTCAACGACAGCTTCCTCGCCCTCAAGGCCGTCCTCCCTCCCGGCTCTAAGAAAGACAAGACATCGATACTGATCAGAGCAAGAGAGTACGTAAAATCTCTCGAGTCAAAGCTGTCAGAGCTGGAGGAGAAGAACCGGGAGCTAGAGGCGAGGCTAGCCAGCCGCCCCGCCGCCGCCGCCAAGAACGACAAAGGCGAGACGGCGGCGGCGCCAGCGCCGGAAGCCGGCGATGAGACGAAGCGAAAGGACCTAGTAGAGATCGAGGTGACGACGAGCGGCGGCGGCGCCGGAGCGGCGGATGCGGCGGCGGCGGCGGGAGGAGATCAAGAGACCTGCACGCTCAACGTAGACTTGCGCGGCGGCGGAGGCGGCGGAGGCATGAGCACGACGGACGTGGTGCTGCGGACGCTGCAGTGCCTGAGAGAGCAGATCGGCGACGGCGCCAGCCTCGTGGCGATGAGCACCAGCGCCGGCTCCGGCGGCCGCCCGCCTCGTGCAAACCTAACATTACAGCTCAAGGTGTAA

>OpDPF

ATGGAGGATTGCAGCAGCTGGAGCCATGGCTACGCCAACGCTACCGGCGGCAACAACGGCTTCATGTGCGGCTACGCTGCCAGCTGCAGCCCCGGAGATCATCAGTTCCAGCAGCAGCAACAGCTGGTCAGCTCGCAGATTCAGCAACACCTCAACGAGATCAACATGCACATGAGCATGGATGACGAGTCGGCGGTGTACGACGGCGCCTCCATGGTGGACGACCTCCTCATGGCTTCCTCGTCGGCGCACCACGCCGGCGCCGGCAGCTTCGTCCAGTACTCCTCCTCGTCCTCCTCTGCCTCCTTCCGCTCCGCCTCCGTCTCATGCAGCCCGGAGAGCTCGGCGGCGGCGACGCACATCCTCGGAGCGCCATCGTCGGCGGCGGCGGGGTTCCATTATTACCCGGAGGTCTCCTCGCAGGCGCCGTTGCCACTACCCTTGCCGCCCTACGAGCCGCAGCACGGCCAATACACCGTCCTCTCGCCGGCGGCGCCAGAGTTGCCGGTGACTACGCCGGCGACCGGCGGCGCGTTCAGGCGCTACGCACGGCACCTCCGCCCGAGGAGGCTGCCCAAGCCGGGAGCTCGCGGGCAGAGGATGTTCAAGACGGCCATGTCGGTGCTCGCCAAGATGCACGTGGCTGCGACGTACAGCCGCCAGTACTACTACCAGCAGGCGGCGGCCGCGTCGGCGTCGGGGGTCGAGGCGGCGGCGCCGCCGTCCGGCAACCAGCTGCAGCACATGATCTCGGAGCGAAAGCGGCGGGAGAAGCTCAACGACAGCTTCCTCGCCCTCAAGACCGTCCTCCCTCCCGGCTCCAAGAAGGACAAGACGTCGATACTGATCAGGGCAAGAGAATACGTAAAGTCTCTCGAGTCAAAGCTGTCAGAGCTGGAGGAGAAGAACCGGAAGCTGGAGGCGCGGTTGGCCAGCCGCCCCGCCGTCGTCGCCGCCAAGAACGACAAAGGCGAGACGGCGGCGGAAGCCGGCGGCGAGACGAAGCGAGAGGACATAGTAGAGATCGAGGTAACGACGAGCGGCACCGGAGCGGCGGATGCTGCGGCGGCGACGGGAGGAGAAGAGACTTGCACGCTCAACGTAGACTTGCGCGGCGGCGGCGGCGGCAGAGGCATGAGCGCGACGGACGTTGTGCTCCGGACGTTGCAGTGCCTGAGAGAGCAGATCGGCGACGGCGCCAGCCTGGTGGCGATGAGCACCAGCGCCGGCTCCGGCGGCCGCCCTCCTCGTGCAAACCTAACATTACAGCTCAAGGTATAA

>OoDPF

ATGGAGGATAGCAGCACCTGGATCCATGGCTACGCCAACGCCAACGCCGCCGGCGGCAACAACGGCTTCATGTGCGACTACGCTGCCAGCTGCAGCCCAGTAGAGtttcagcagcagcaacagctggTCAGCTCGCAGATTCAGCTACACCTCAACCAGATCAACATGCACATGAGCATGGATGACGAGTCGGCGGTGTACGACGGCGCCTCCATGGTGGACGACCTCCTCATGGCTTCCTCGTCGGCGCAccacgccggcgccggcagCTTCCAGTactcctcgtcgtcctcctctgcCTCCTTCCGCTCCGCCTCCGTCTCATGCAGCCCGGAGAGCTCGGCAACGGCGACGCACATCCTCGGAGCCCcagcgtcggcggcggtggggttCCATTACCCGGAGGTCTCCTCGCAGGCGCCGTTGCCACTACCCTTGCTGCCCTACGAGCCGCAGCACGGCCAATACACCGTCCTCTCGCGGCCGGCGCCAGAGTTGCCGGCGACTACGCCGGCGACCGGCGTGTTCAGGCGCTACGCGCGGCACCTCCGCCCGAGGAAGCTGCCCAAGCCGGGAGCTTGCGGGCAGAGGATGTTCAAGACGGCCATGTCAGTGCTCGCCAAGATGCACGTGGCGGCGACGTACAGAAGCCAGTACTACTACCAGCAGGCGGCAGCCACGTCGGCTTcggcggccgaggcggcgccgccgccgtccggcAACCAGCTCCAGCACATGATCTCGGAGCGCAAGCGGCGGGAGAAGCTCAACGACAGCTTCCTCGCCCTCAAGGCCGTCCTCCCTCCCGGCTCCAAGAAAGACAAGACGTCGATACTGATAAGAGCAAGAGAGTACGTAAAGTCTCTCGAGTCAAAGCTGtcagagctggaggagaagaaccgggagctggaggcgcggTTGGCCAGCCgccccgccgtcgtcgccgccaagAACGACGAAGGCGAGAtagcgacggcggcgccggaaGCCGGCGACGAGACGAAGCGAGAGGACCTAGTAGAGATCGAGGTgacgacgagcggcggcggcgccggagcggcggcgacgggaggaGAAGAGACTTGCACGCTCAACGTAGActtgcgcggcggcggcggcggcggaggcatgAGCACGACGGACGTTGTGCTCCGGACGCTGCAGTGCCTGAGAGAGCAGATCGGCGACGGCGCCAGCCTGGTGGCGATGAGTACCAGTGCCGGCTCCGGCGGCCGCCCTCCTCGTGCAAACCTAACATTACAGCTAAAGGTATAA

>ObDPF

ATggatagcagcagcagcagctggatCCATGGctacgccggcgccggcgccggcggcaacaACGGCTTCATGTGCGGCTACGCTGCCAGCTGCAACCCCGGTGAGtttgagcagcagcaggtggtgGTCAGCTCGCAGATTCAGGACCAGCTCAACCAGATCAGCATGCACATGAGCAtggacgacgagtcggcggtGTACGACGGCGCCTCCATGGTGGAGGTGGACGACCTGCTGATGACCTCGTCGGCGCACCACGGCGGCTgcttcacctcctcctcgtcgtcgtctgcttccttcccctccgcctccgtctCATGCAGCCcggagagctcggcggcggcgcacgtcCTCGGagcgccagcggcggcggccgcggggtTCCTGTACCCGGAGGTCTCGTCGCAGGCGGCGCCGTTGCCACTGCGACGGCCACCCTTGGTGCCATACGAGCCCGAGCCCCAGCACCAGTGCCACGGCAGCTACACCGGCCTGcagtcgccggcgagcggccgcggcgcgtTCAAGCGGTACGCGCGGCACCTCGGCCCGAGGCGGCCGCCCAATAAGGCGCCCGGGCAGAGGATGTTCAAGACGGCCATGTCGGTGCTGGCCAAGATGCACGTGGCGATGACGTACAGCCGGCAGTACTACTaccagcaggcggcggcggcggaggcggaggcgccggagccgccgtCCGGCAACCAGCTGCAGCACATGATCTCCGAGCGCAAGCGGCGGGAGAAGCTCAACGACAGCTTCGTCGCCCTCAAGGCCGTCCTCCCTCCCGGCTCCAAAAAAGACAAGACGTCGATACTGATCAGGGCAAGGGAGTACGTGAAGTCACTCGAGTCCAAGCTGTCGGAGCTGGAGAAGAAGAACCGGGAGCTGGAGGCGCGGCTGGCCATGGCCAAGAACGACGGgagcgagacggcggcggcagcggcggcggcacccgacgccgccggcgacgacacgaAGCGAGAGGACGAACTAGTAGAGATCGAGCtgaccagcggcggcgccggagcggCGGTGGGATCAGGCCAAGACGAGGAGACCTGCTGCACGCTCAACGTAGCCGTGACGCCACCGCCatcgcgacgcggcggcggcggcggcatgagCACGACGGACGTCGTGCTCCGGACGCTGCAGTGCCTGAGGGAGCagatcggcgacggcgccagcCTGGTGGCGATGAGCACCagcgccggctccggcggccgccCTCCTCGAGCGAACCTAACATTACAGCTTAaggtataa

>LpDPF

ATGGAGAGCAGCAGCTGGATCCATGGCGGCTACGCCCACGCCAACGGCGCCAGCAACAACGGATTCATGTGCGGCTATGCTTCCAGTTGCAGCCCTGGGGAATTTCAGCTCAAGGAGCAGCAGCAACAGCAACAACTAGTCAGCTCTCAGATTCAGCACCATCTCAACCAGATTAGCATGCACATGAGCATGGATGACGATCAGTCAACGGTCTACGACGGCGCCGCCATGGACGACCTCTTCATTCCTTCCGGCAGCTTCCCTTCCTCCTCCTCTTCCTCCGCCTCCTACCGCTCCACCTCCGTCTCCTACAGCGCCGACACCTCGCCGGCGACGGCGGCGCCGCACGTCCTCGGAGCACCGGCGCCGGCGGCCGGATTCATCCAATTCCCGGAGGTCTCCTCGCACGCGCCGCCGTACACCGGGTTCTCGCCGCCGGCTGCTAGCGGCGGCGCGTTCCGGCGATACGCGCGCCACCTGGGGCCGAGGAGGGCTGTGGCGAAGAGCGGCGGAGGGCAGAGGATGTTCAAGACGGCGATGTCGGTGCTGTCGAAGATGCACGTGGCGGCGATGGCGTATAGCCGGCAGTGCTACTACCAGCAGCAGCAGGCGGCAGCGGCGGCGGCGGAGGCGGCGCCGTCCGGCAACCAGCTGCAGCACATGATCTCCGAGCGCAAGCGGCGTGAGAAGCTCAACGACAGCTTCGTCGCCCTCAAATCCGTCCTCCCTCCCGGCTCCAAGAAAGACAAGACGTCGATACTGATCAGAGCAAGAGAATACGTGAAGTCACTCGAGTCAAAGCTATCGGAGCTGGAGGAGAAGAACCGGGAGCTGGAGGCGCGGCTAGCCTCGACGGCGGCGCCACCGGCCGCCGCCGCCGCCACCGTTGAGAAGGGAGAAGAAATAGTAGAAATCGAGGTGACCAGCGCCGGCGCCGGCGCCGGCGCCGGAGCAAATGGAGAAGATCAGGAGAGCACGTGCACACTGAACGTGGCGGTGACGACGCCGCCGGCGACGTCGCGCGGCGGCGGCGGCGGAATGAGCACGACGGACGTGGTGCTCCGTACGCTGCAGTGTCTGAGAGAGCAGATCGGTGACGGAGCAAGCTTGGTGGCGATGAGCACCAGCGCCGGCTCCGGCGGCCGGCCGCCTCGAGCAAACCTAACATTACAGCTTAAGGTATAA
